# BrainSuite BIDS App: Containerized Workflows for MRI Analysis

**DOI:** 10.1101/2023.03.14.532686

**Authors:** Yeun Kim, Anand A. Joshi, Soyoung Choi, Shantanu H. Joshi, Chitresh Bhushan, Divya Varadarajan, Justin P. Haldar, Richard M. Leahy, David W. Shattuck

## Abstract

There has been a concerted effort by the neuroimaging community to establish standards for computational methods for data analysis that promote reproducibility and portability. In particular, the Brain Imaging Data Structure (BIDS) specifies a standard for storing imaging data, and the related BIDS App methodology provides a standard for implementing containerized processing environments that include all necessary dependencies to process BIDS datasets using image processing workflows. We present the BrainSuite BIDS App, which encapsulates the core MRI processing functionality of BrainSuite within the BIDS App framework. Specifically, the BrainSuite BIDS App implements a participant-level workflow comprising three pipelines and a corresponding set of group-level analysis workflows for processing the participant-level outputs. The Anatomical Pipeline extracts cortical surface models from a T1-weighted (T1w) MRI. It then performs surface-constrained volumetric registration to align the T1w MRI to a labeled anatomical atlas, which is used to delineate anatomical regions of interest in the MRI brain volume and on the cortical surface models. The Diffusion Pipeline processes diffusion-weighted imaging (DWI) data, with steps that include coregistering the DWI data to the T1w scan, correcting for susceptibility-induced geometric image distortion, and fitting diffusion models to the DWI data. The Functional Pipeline performs fMRI processing using a combination of FSL, AFNI, and BrainSuite tools. It coregisters the fMRI data to the T1w image, then transforms the data to the anatomical atlas space and to the Human Connectome Project’s grayordinate space. The outputs of each pipeline can then be processed during group-level analysis. The outputs of the Anatomical Pipeline and the Diffusion Pipeline are analyzed using the BrainSuite Statistics Toolbox in R (bstr), which provides functionality for hypothesis testing and statistical modeling. The outputs of the Functional Pipeline can be analyzed using atlas-based or atlas-free statistical methods during group-level processing. These analyses include the application of BrainSync, which synchronizes the time-series data temporally and enables comparison of resting-state or task-based fMRI data across scans. We also present the BrainSuite Dashboard quality control system, which provides a browser-based interface for reviewing the outputs of individual modules of the participant-level pipelines across a study in real-time as they are generated. BrainSuite Dashboard facilitates rapid review of intermediate results, enabling users to identify processing errors and make adjustments to processing parameters if necessary. The comprehensive functionality included in the BrainSuite BIDS App provides a mechanism for rapidly deploying the BrainSuite workflows into new environments to perform large-scale studies. We demonstrate the capabilities of the BrainSuite BIDS App using structural, diffusion, and functional MRI data from the Amsterdam Open MRI Collection’s Population Imaging of Psychology dataset.

## 1 Introduction

The neuroimaging community has made considerable progress in recent years to establish new methods and standards that promote open and reproducible science. These efforts have included the recommendation of best practices for analyzing and sharing data (Nichols et al., 2017; Pernet et al., 2020), the development of the Brain Imaging Data Structure (BIDS) to standardize data organization for brain imaging studies (K. J. Gorgolewski et al., 2016), and the development of the BIDS App methodology (K. J. Gorgolewski et al., 2017) for improved portability and reproducibility. BIDS Apps encapsulate analysis workflows within software containers, which can process study data organized according to the BIDS standard. The BIDS App framework addresses an important challenge in neuroimaging analysis, which is to ensure that the processing sequences that are applied to data are maintained such that they can be easily shared and reproduced. A BIDS App container includes all software and reference data necessary to apply a particular image processing workflow to a new dataset. It ensures that consistent versions of the operating system, software libraries, binaries, scripts, and reference data are provided for a specific analysis sequence.

Conventional distribution of image processing pipelines presents several potential issues related to reproducibility, portability, and interoperability. In the context of software, reproducibility requires that programs can be rerun identically on a set of data to produce the same result. Reproducibility is essential to ensure that study data are processed consistently to avoid introducing bias, particularly in the case of multi-site or longitudinal studies (Jovicich et al., 2009; Kruggel et al., 2010). In general, reproducibility in neuroimaging is a challenging issue due to the wide variety of advanced and complex image processing and analysis methods, which often integrate programs and data from multiple software packages developed by different research groups.

Numerous issues can arise when these workflows are shared, including: 1) the need for users to install software packages and their dependencies, possibly in environments where they require escalated privileges or assistance from system administrators; 2) execution on platforms upon which the workflows were not tested; 3) execution in environments that have different versions of the software components than were used during development; 4) upgrades in versions of software programs or support libraries during routine maintenance, which possibly occur without the user’s knowledge; 5) adjustments to parameter settings in individual software components, which may not be well-documented; and 6) variations in required input data format and organization across different software programs. These issues can lead to difficulties and burdens for end users in terms of installation and data organization, as well as potentially producing statistically significant differences when the same workflow is used on the same data in different computational environments (Glatard et al., 2015; Gronenschild et al., 2012).

The BIDS App framework was created to address these issues by providing a mechanism for the development and archival of portable applications using lightweight container technologies (K. J. Gorgolewski et al., 2017). All BIDS Apps must meet three basic requirements. The first is that they must be compatible with datasets that follow the BIDS specification, which provides standards for file structures, naming conventions, and data descriptions for neuroimaging data (K. J. Gorgolewski et al., 2016). BIDS requires specific file formats, including the Neuroimaging Informatics Technology Initiative (NIfTI), JavaScript Object Notation (JSON), and tab separated value (TSV) formats. By using standardized data formats, BIDS improves interoperability and facilitates the use of previously acquired datasets. Because BIDS Apps require input data to be organized, named, and formatted in a consistent manner, users can run different BIDS Apps on the same dataset without renaming or reorganizing the files. Second, BIDS Apps must be packaged in a Docker container (Merkel, 2014) that can be converted into an Apptainer (formerly known as Singularity) image (Kurtzer et al., 2017). The availability of an Apptainer image ensures that users have a version of the BIDS App that is more secure and that is suitable for use on multi-user systems with shared resources. Third, BIDS Apps must implement a command-line interface that uses a common set of arguments. Each BIDS App is required to accept three core positional arguments: 1) input dataset directory, 2) output directory, and 3) stage of analysis. The third argument specifies either participant-level or group-level processing. This division enables participant-level processing to be executed in parallel if appropriate compute resources are available. The containerization of the software simplifies its deployment on a cluster environment, thus facilitating this parallelization. There have been several BIDS Apps released over the past few years, including more than 40 that are currently maintained in the BIDS App Repository (https://bids-apps.neuroimaging.io/) hosted by the Center for Reproducible Neuroscience (CRN) at Stanford University (please see Discussion for a description of some of the existing BIDS Apps).

In this paper, we present the BrainSuite BIDS App, which encapsulates components of BrainSuite into the BIDS App framework. BrainSuite is a collection of open-source software tools for analyzing brain imaging data, which we have been developing for more than two decades (e.g., Bhushan et al., 2015, 2016; Haldar & Leahy, 2013; A. A. Joshi et al., 2007; A. A. Joshi, Bhushan, et al., 2018; A. A. Joshi, Chong, et al., 2018; A. A. Joshi et al., 2012, 2022; J. Li et al., 2018; Shattuck & Leahy, 2002; Shattuck et al., 2001; Varadarajan & Haldar, 2018). The BrainSuite BIDS App implements a comprehensive workflow comprising three pipelines to analyze anatomical T1-weighted (T1w) MRI, diffusion MRI (dMRI), and functional MRI (fMRI) data. We note that these three BrainSuite BIDS App pipelines each make use of BrainSuite command-line programs that can be configured with greater flexibility in their stand-alone forms. These command-line programs compose the existing stand-alone BrainSuite pipelines, named the BrainSuite Anatomical Pipeline, the BrainSuite Diffusion Pipeline, and the BrainSuite Functional Pipeline.

In the present work, we have selected a subset of the available pipeline and program options for use in the BrainSuite BIDS App to integrate their different components into a cohesive workflow. The functionality that has been incorporated from each of the BrainSuite standalone pipelines into the BrainSuite BIDS App is as follows. The BrainSuite Anatomical Pipeline (BAP) processes T1w MRI to perform tissue classification and extract cortical surface models (Shattuck & Leahy, 2002; Shattuck et al., 2001), to estimate subject-level measures of cortical thickness (A. A. Joshi, Bhushan, et al., 2018), to align cortical surface and volume data to a labeled brain atlas (A. A. Joshi et al., 2022) using surface-constrained volumetric registration (A. A. Joshi et al., 2007; A. A. Joshi et al., 2012), and to estimate various neuroanatomical measures. The BrainSuite Diffusion Pipeline (BDP; Varadarajan et al., 2020) performs susceptibility-induced geometric image distortion correction and registration of the diffusion MRI data to the anatomical T1w image (Bhushan et al., 2012, 2015), fits diffusion tensor models to the distortion-corrected diffusion MRI, and generates maps of diffusion parameters (e.g., fractional anisotropy, mean diffusivity). BDP also provides a selection of methods for estimating orientation distribution functions for more complicated diffusion sampling schemes (e.g., Haldar & Leahy, 2013; Varadarajan & Haldar, 2018). The BrainSuite Functional Pipeline (BFP) provides methods for analyzing resting-state and task-based fMRI data using a combination of tools from BrainSuite (Bhushan et al., 2016; J. Li et al., 2018), FSL (S. M. Smith et al., 2004) and AFNI (Cox, 1996). Processed fMRI data are transformed into the grayordinate representation defined by the 32K Conte-69 surface, a standard surface-volumetric coordinate system developed for the Human Connectome Project (Glasser et al., 2013).

The BrainSuite BIDS App makes use of the stand-alone BrainSuite pipelines and augments them with additional steps to facilitate group-level analysis and to perform some additional functionality (see Sec. 2.4 for details). We refer to the BrainSuite BIDS App versions of the pipelines in this paper simply as the Anatomical Pipeline, the Diffusion Pipeline, and the Functional Pipeline to distinguish them from the existing standalone versions, which we reference using their established names and acronyms (BrainSuite Anatomical Pipeline [BAP], BrainSuite Diffusion Pipeline [BDP], and BrainSuite Functional Pipeline [BFP]).

Each participant-level BrainSuite BIDS App pipeline has a corresponding group-level analysis component. The outputs of the Anatomical Pipeline (e.g., cortical thickness or regional volume measures) can be compared across subjects or time points during the group-level analysis stage using the BrainSuite Statistics Toolbox in R (bstr; S. Joshi et al., 2020, 2023). These comparisons are made in the common atlas space. For group-level analysis of the Diffusion Pipeline outputs, diffusion parameter maps are resampled from the subject T1w space in which they were computed to the atlas space, where group-level comparisons can be performed using bstr. The Functional Pipeline outputs are compared at the group level after applying BrainSync (A. A. Joshi, Chong, et al., 2018), which synchronizes resting-state fMRI data temporally between the subject dataset and a reference dataset. The BFP program can perform either atlas-based analysis using a reference dataset created from multiple input datasets (Akrami et al., 2019) or atlas-free statistical testing where pair-wise comparisons of all pairs of subjects is performed and used as test statistics for regression or group difference studies (A. A. Joshi et al., 2021).

The BrainSuite BIDS App facilitates distributed processing of data, enabling large numbers of subjects to be processed concurrently in cluster computing environments. As dataset sizes increase, this also increases the burden on users to evaluate the generated outputs and identify errors in processing that may require intervention. To address this issue, we developed BrainSuite Dashboard, a browser-based quality control system for rapidly evaluating the outputs of the BrainSuite workflows. This system generates snapshot images for key stages in the participant-level workflows, which are displayed in an interactive web page that is updated in real time while a set of BrainSuite BIDS App instances process a study dataset. This can assist the user in determining if any of the data need to be reprocessed, if settings need to be modified, or if data need to be excluded from a study.

The BrainSuite BIDS App is freely available as a buildable Docker script, a pre-built Docker image, and a pre-built Apptainer image. The BIDS App mechanism provides users with the ability to apply the primary BrainSuite image analysis routines with minimal installation effort, as well as to rapidly evaluate intermediate outputs at the participant level. In the following sections, we describe the BrainSuite BIDS App in greater detail and demonstrate its functionality and utility by applying it to the Population Imaging of Psychology dataset from the Amsterdam Open MRI Collection (Snoek et al., 2021).

## 2 Methods

### 2.1 Architecture

Following the BIDS App methodology, we implemented the BrainSuite BIDS App with a participant-level workflow and a group-level analysis workflow, which are integrated as illustrated in Fig. 1. The BrainSuite BIDS App container includes complete R and Python environments, as well as all necessary packages and dependencies to perform the participant-level and group-level analyses. This eliminates the need for the user to install and configure these, maintains a consistent set of packages, and does not interfere with the user’s local environment on the host machine. Each workflow component can be configured using a set of JSON files (see App. A.1 for details). These settings can be specified either at the study level for all subjects or at the participant level if individual configuration files are included. The participant-level workflow, shown in detail in Fig. 3, includes the Anatomical Pipeline, the Diffusion Pipeline, and the Functional Pipeline. These make use of components from the corresponding stand-alone BrainSuite pipelines. By default, each pipeline will be run if the data it requires are present in the BIDS dataset. The user can also configure the BrainSuite BIDS App to omit individual pipelines (see App. A.1). The intermediate outputs of the participant-level workflows can be readily visualized using our browser-based BrainSuite Dashboard, which provides live feedback to the user to assist in the detection of processing errors or other issues. The group-level workflow uses the BrainSuite Statistics Toolbox in R (bstr) to analyze the outputs of the Anatomical Pipeline and Diffusion Pipeline, and it uses a set of Python-based tools to analyze the outputs of the Functional Pipeline.

**Figure 1:**
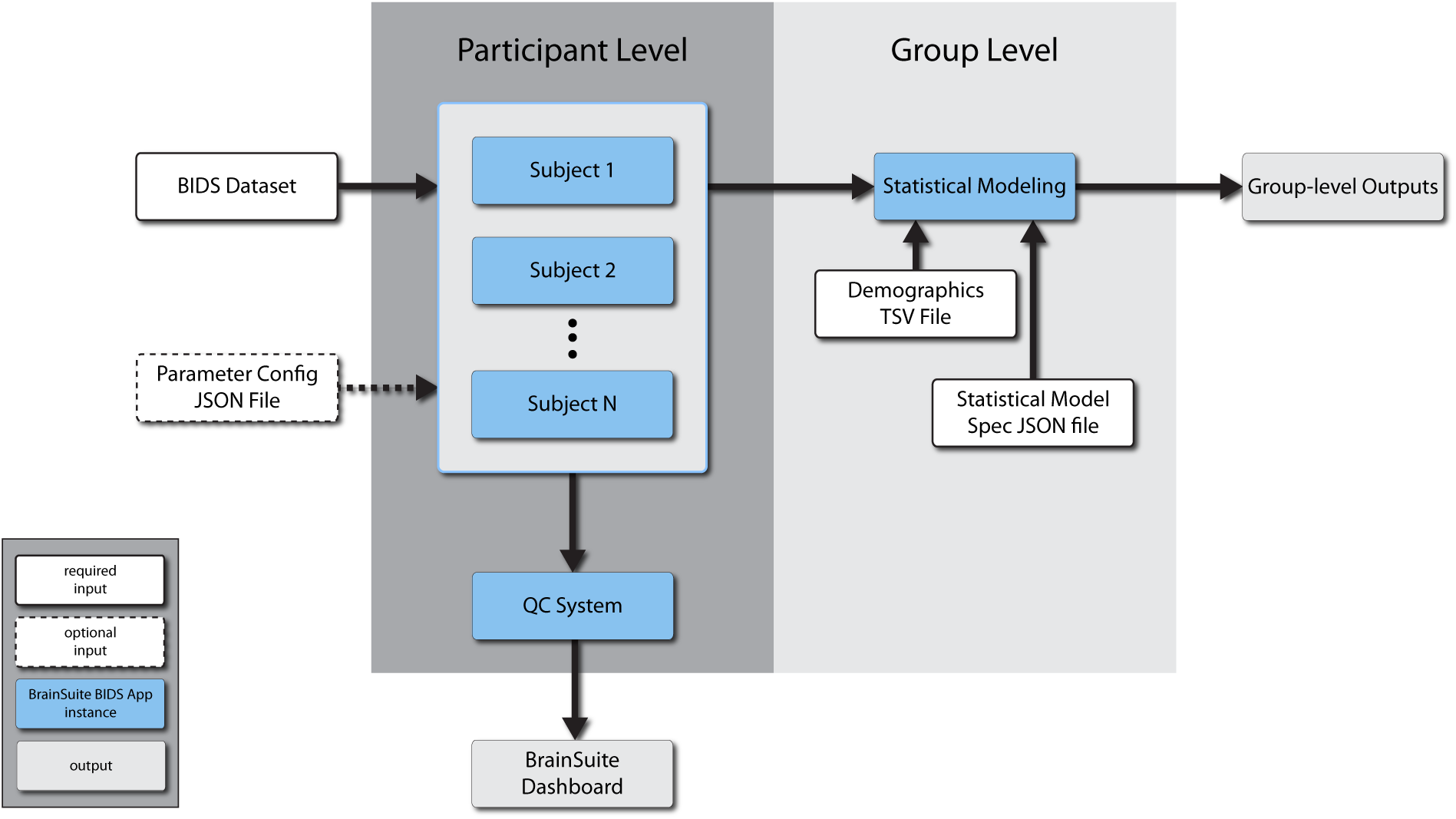
BrainSuite BIDS App Architecture. The BrainSuite BIDS App performs complete analysis sequences for structural, functional, and diffusion MRI data stored in the BIDS format (see Fig. 2). At the participant level, data for each subject can be processed in a separate BrainSuite BIDS App instance, which can be run concurrently. These execute the participant-level workflows (see Fig. 3). The specific processing performed by the participant-level workflows can be controlled by an optional configuration JSON file, which specifies parameters for the different stages of processing (see A.1). A separate BrainSuite BIDS App instance can be invoked to run as the Quality Control (QC) System, which monitors the outputs of the individual participant-level instances and generates process snapshots and status codes. These are read by the BrainSuite Dashboard system (see Fig. 4), which provides a browser-based dynamic view of the participant-level results as they are generated. The outputs of the participant-level instances are processed by a group-level instance, which performs statistical testing specified by a statistical model JSON file based on participant demographics stored in a TSV file. Table 1 details the participant-level output measures and corresponding group-level analyses available in the BrainSuite BIDS App.

**Table 1:** BrainSuite BIDS App Outputs. Output measures generated at the participant levels, and the corresponding group-level analyses available for each type of data.

|  | Participant-level Measures | Group-level Analyses* |
| --- | --- | --- |
| Anatomical | cortical surface meshes (inner, pial, mid) | thickness maps in atlas surface space |
|  | GM, WM, CSF tissue fractions (per voxel) | analysis of gross GM, WM, CSF volume measures |
|  | thickness and surface area for cortical ROIs | ROI-based analysis of surface area and mean thickness cortical measures |
|  | GM and WM volume within each ROI | ROI-based analysis of GM and WM volume measures |
|  | volume Jacobian determinants | relative volume differences from Jacobian maps |
| Diffusion | diffusion parameter maps (FA, MD, AxD, RD, GFA) | voxel-wise analysis of FA, MD, AxD, RD, GFA |
| Functional | task or resting fMRI data in grayordinate space after filtering and motion correction | statistical maps in grayordinate space in surface and volume space |

\*Available statistical tests for anatomical and diffusion analysis include ANOVA, correlation, t-test, and paired t-test. Available statistical tests for functional MRI analysis include atlas-based similarity test and atlas-free pairwise tests.

The BrainSuite BIDS App’s Anatomical Pipeline, Diffusion Pipeline, and Functional Pipeline are each composed of several executable programs, most of which are compiled C++ or MATLAB code. The Anatomical Pipeline uses programs that are all included in the BrainSuite distribution. The Diffusion Pipeline relies primarily upon the stand-alone BrainSuite Diffusion Pipeline (BDP) executable. In the BrainSuite BIDS App, we have augmented this with FSL’s eddy (Andersson & Sotiropoulos, 2016), which is installed using the FSL-conda repository (https://fsl.fmrib.ox.ac.uk/fsldownloads/fslconda/public/). The Functional Pipeline is built upon the BrainSuite Functional Pipeline (BFP). BFP also employs components from AFNI (Cox, 1996) and FSL (S. M. Smith et al., 2004), which are imported in the BrainSuite BIDS App via NeuroDebian repositories (Y. O. Halchenko & Hanke, 2012). These individual components are connected within the BrainSuite BIDS App using Nipype (K. Gorgolewski et al., 2011), a Python-based API that enables users to execute various programs using a common interface. Nipype provides Python objects that wrap programs and tools from a heterogeneous collection of APIs. These objects, termed nodes, can be assembled to create workflows by connecting their inputs and outputs. The Nipype pipeline can then be executed, with individual modules triggering subsequent processing steps upon their completion. This controls the workflow by running processes when resources become available. Processes that are scheduled to run in parallel will begin if compute resources are available. This job scheduling prevents processes from contending for resources and enables computationally efficient processing.

We created a set of Nipype Python objects to provide interfaces to the command-line tools included in the BrainSuite 23a distribution. We then developed a Python script that connects instances of these objects to create the Nipype pipelines. The BrainSuite BIDS App’s Diffusion Pipeline and Functional Pipeline both depend on outputs from its Anatomical Pipeline. These data dependencies are managed by Nipype, which launches the Diffusion Pipeline and the Functional Pipeline when their required inputs have been generated (see Fig. 3). The Diffusion Pipeline requires the output of the nonuniformity correction stage of the Anatomical Pipeline; upon completion of this stage, Nipype triggers the start of the Diffusion Pipeline, which runs in parallel with the remaining steps of the Anatomical Pipeline. The Anatomical Pipeline will wait for the completion of the atlas-registration stage before performing the resampling of DTI parameter maps to the atlas space. Similarly, when the Anatomical Pipeline completes the registration and labeling of the T1w and cortical surface, Nipype will start the Functional Pipeline. The Functional Pipeline will run concurrently with the Diffusion Pipeline if the Diffusion Pipeline has not yet finished. If multiple fMRI datasets are included for processing, the Functional Pipeline will be applied to each dataset serially.

### 2.2 Integration, Testing, and Deployment

Throughout the development of the BrainSuite BIDS App, we used CircleCI to perform continuous integration and to conduct a series of tests each time code changes were committed to our Git repository. We designed these tests to ensure that the BrainSuite BIDS App code will run successfully when launched and that it will also meet Apptainer’s read-only criterion. We submitted the BrainSuite BIDS App to the BIDS App Repository (https://bids-apps.neuroimaging.io/), where it underwent the standard set of validation tests applied to all BIDS Apps to ensure that they meet architectural and security requirements. After successfully passing these tests, the BrainSuite BIDS App was approved for distribution and made available at https://github.com/BIDS-Apps/BrainSuite, along with user documentation written in markdown files. Any changes to the release version of the BrainSuite BIDS App or the markdown files are archived in the CRN BIDS Apps GitHub repository. We also released a pre-built Docker image on DockerHub (https://hub.docker.com/r/bids/brainsuite) as well as a pre-built Apptainer image at https://brainsuite.org/data/sif/bids_brainsuite_v23a.simg. The user can optionally convert the Docker-based BrainSuite BIDS App into an Apptainer image, which requires approximately 6 minutes on a computer with 4 cores and 64GB of RAM.

### 2.3 BrainSuite BIDS App Inputs

The BrainSuite BIDS App assumes that the input data are stored according to BIDS conventions. For some users, this may require conversion of study data into NIfTI (e.g., from DICOM), generation of JSON files that include descriptive information regarding the study data, and copying or moving of data into the BIDS directory hierarchy. There are several freely available BIDS converters, including BIDScoin (Zwiers et al., 2022), heudiconv (Y. Halchenko et al., 2022) and Dcm2Bids (Bedetti et al., 2022). A basic example of a BIDS data directory is shown in Fig. 2. All data for a study are stored within one top-level directory, which contains a JSON file describing the dataset and a set of sub-directories for each subject in the study. Within each subject directory are sub-directories for anatomical, diffusion, and functional MRI. Additional modalities are also allowed in BIDS, but the BrainSuite BIDS App will not process these. In addition to the layout shown in Fig. 2, each subject may have data from multiple sessions, which would be stored in sub-directories immediately below the subject directory, with the session directories containing anat, dwi, and func directories as necessary (see the BIDS specification (BIDS-Contributors, 2022) for details). An anat directory containing at least one T1-weighted MRI is required for BrainSuite BIDS App to run. Diffusion and functional data will be processed if present in the dwi and func directories, respectively.

**Figure 2:**
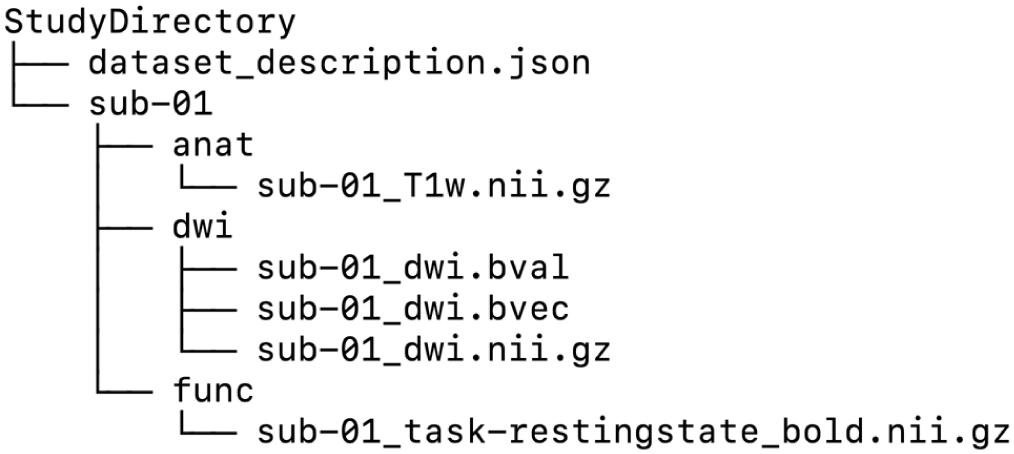
Example BIDS data directory. Shown is an example of data from one subject organized in a BIDS-Compliant directory structure following the BIDS file naming scheme. Only the anatomical T1w MRI is required to run the BrainSuite BIDS App.

### 2.4 Participant-level Workflow

As detailed in Fig. 3, the participant-level workflow comprises three primary BrainSuite BIDS App pipelines. Each component can be configured to be run separately. The BrainSuite BIDS App’s Diffusion Pipeline and Functional Pipeline both require outputs from its Anatomical Pipeline to run, though these outputs may have been computed previously in a separate BrainSuite BIDS App session or outside of the BrainSuite BIDS App. The implementations of these BrainSuite BIDS App pipelines mirror the corresponding pipelines included in the BrainSuite 23a distribution, with a few modifications that facilitate integration of their functionality into a comprehensive subject-level workflow. The BrainSuite BIDS App’s Diffusion Pipeline includes an optional preprocessing stage for eddy current and motion correction using FSL’s eddy (Andersson & Sotiropoulos, 2016), which is not included in the stand-alone versions of BDP. We have also implemented additional workflow components that are designed to prepare the participant-level data for use in the group-level analysis stage. In the BrainSuite BIDS App’s Anatomical Pipeline and Diffusion Pipeline, we have added stages that perform image- and surface-based smoothing to the stand-alone versions of these pipelines. The stand-alone BrainSuite Functional Pipeline (BFP) distribution includes participant-level and group-level processing. We use the BFP participant-level component in the BrainSuite BIDS App Functional Pipeline. The BFP tools for synchronizing and analyzing fMRI data at the group level are included in the BrainSuite BIDS App Group-level Analysis stage. Importantly, many of the tools and pipelines used in the BrainSuite BIDS App have greater flexibility when invoked separately on the command line or within the BrainSuite GUI (more details of each component’s full functionality are provided on the BrainSuite website). As described in App. A.1, we have implemented the capability to customize several program options through the use of JSON-formatted configuration files. As the BrainSuite BIDS App evolves and new use cases emerge, we anticipate expanding the configuration options to include additional features and new functionality. We note that additional preprocessing steps can also be performed prior to running the BrainSuite BIDS App, provided that the data are placed in the input directory according to the BIDS specifications. In the following sections, we assume that all necessary input data for each pipeline are in the BIDS directory and that each pipeline has been configured to be executed.

**Figure 3:**
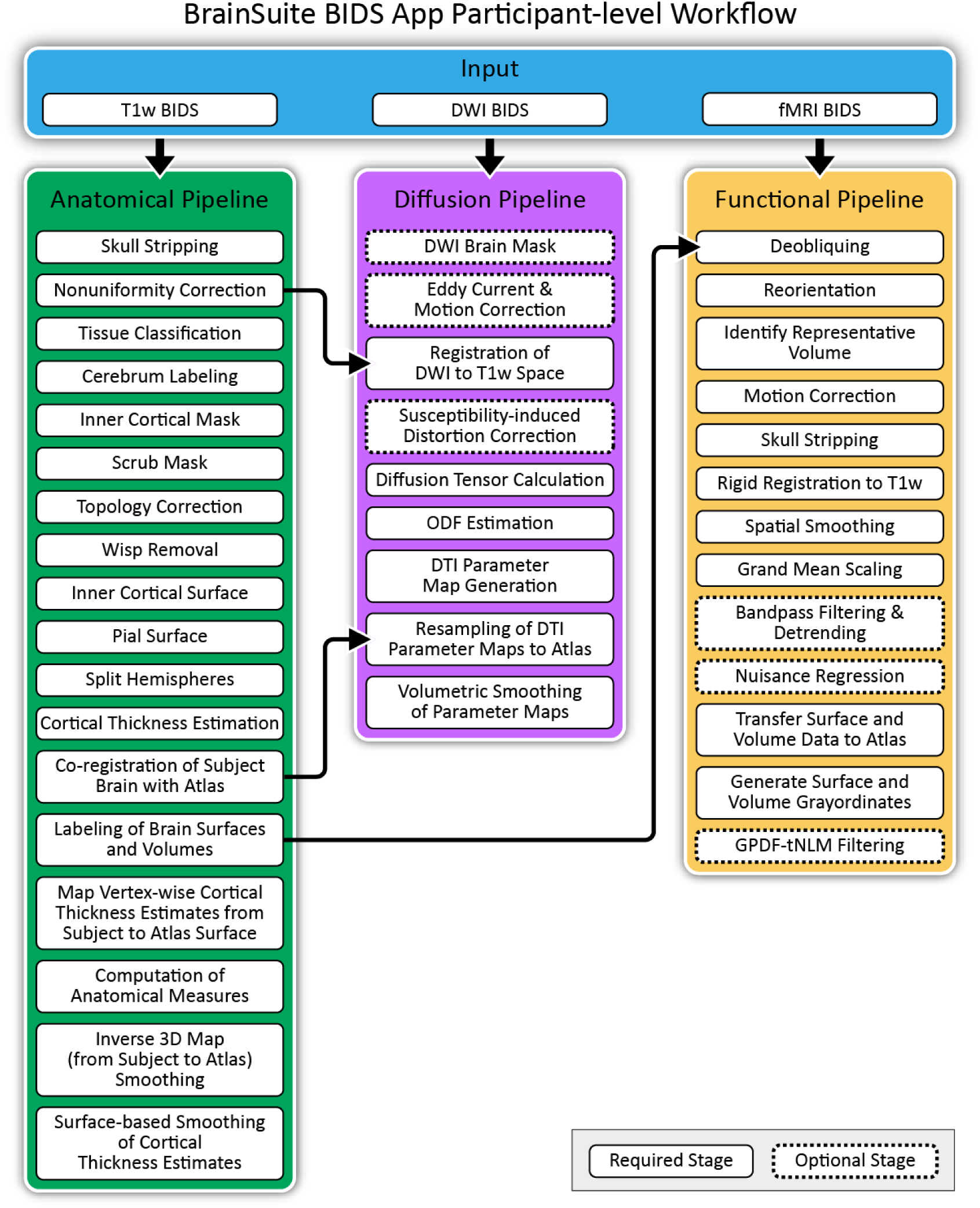
BrainSuite BIDS App Participant-level Workflow. Shown is the BrainSuite BIDS App participant-level workflow, including the stages of the Anatomical, Diffusion, and Functional Pipelines. An anatomical T1-weighted MRI is required. Diffusion MRI and functional MRI data will also be processed if present in the BIDS directory hierarchy. The Diffusion Pipeline performs registration-based susceptibility-induced geometric image distortion correction using the nonuniformity-corrected skull-stripped image output by the second stage of the Anatomical Pipeline. The Anatomical Pipeline creates mappings from the subject’s anatomical T1w image to an atlas space. These mappings are used by the Diffusion Pipeline and the Functional Pipeline to map the diffusion and functional data into the atlas space for group-level studies.

#### 2.4.1 Anatomical Pipeline

The BrainSuite BIDS App Anatomical Pipeline, shown in green in Fig. 3, takes a NIfTI file containing a T1-weighted brain MRI as input. The Anatomical Pipeline first generates surface models of the inner and outer boundaries of the cortical surface using a series of steps that include skull and scalp removal, nonuniformity correction, tissue classification, cerebrum identification, topology correction, and surface generation (Shattuck & Leahy, 2002; Shattuck et al., 2001). The Anatomical Pipeline estimates cortical thickness using a method based on the anisotropic diffusion equation (ADE), which uses the tissue fraction maps generated during tissue classification (A. A. Joshi, Bhushan, et al., 2018). The output of this stage is an individual subject surface mesh that includes cortical thickness estimates at every vertex of the mesh.

The next stage of the Anatomical Pipeline registers the subject surface and volume data to a labeled atlas using surface-constrained volumetric registration (SVReg; A. A. Joshi et al., 2007; A. A. Joshi et al., 2012). SVReg first performs surface-based registration to match the subject cortical surface mesh to the atlas mesh. It then performs volumetric registration to the image volume of same atlas, while constraining the coregistered cortical surfaces to remain fixed. This produces consistent surface and volume mappings between the individual subject and the atlas, which enables data to be mapped between these spaces. SVReg then uses these mappings to transfer labels, which define anatomical regions of interest (ROIs), between the atlas and the subject. This results in a labeling of neuroanatomical structures that is consistent between the surface and the volume. The BrainSuite BIDS App includes three labeled brain atlases: BrainSuiteAtlas1, which includes 99 anatomical ROIs defined on the Colin27 atlas (Holmes et al., 1998); the BCI-DNI Brain Atlas, which includes 95 ROIs defined on a high-resolution single subject atlas; and the USC Brain Atlas, which includes 159 ROIs defined by functional subdivisions of the BCI-DNI Brain Atlas (A. A. Joshi et al., 2022). The user can specify the atlas selection as a parameter when invoking the BIDS App. The Anatomical Pipeline then uses the subject-to-atlas surface mapping generated by SVReg to map the cortical thickness estimates from the subject surface space to the atlas surface space. This step prepares the data for group comparison of surface data. The Anatomical Pipeline also computes a series of subject-specific scalar metrics, including measures of surface area, volume, and average grey matter thickness of each anatomical ROI.

The Anatomical Pipeline produces multiple outputs that can be analyzed using bstr during the group-level analysis stage. These include labeled cortical surfaces, labeled image volumes, the deformation fields that map from the atlas to each subject, and the corresponding inverse deformation fields (see Table 1). In preparation for group-level analysis, the Anatomical Pipeline performs spatial smoothing of the surface and volume data to be analyzed. The purpose of these smoothing operations is to increase the signal-to-noise ratio and to compensate for mismatch due to anatomical variation and registration errors. For tensor-based morphometry (TBM), the Anatomical Pipeline applies isotropic Gaussian smoothing to the Jacobian determinants computed from the inverse deformation maps produced by atlas-to-subject registration. By default, the Gaussian smoothing uses a kernel of size *σ* = 3 mm; this is configurable by the user. For cortical surface thickness data, the thickness estimates at each vertex are smoothed using the Laplace-Beltrami operator (A. A. Joshi et al., 2009). The default smoothing level for surface data is *σ* = 2 mm, but this is also modifiable by the user. We note that these smoothing steps are not part of the command-line or GUI versions of the BrainSuite Anatomical Pipeline, but they can be run separately using command-line programs that we provide in the BrainSuite distribution.

#### 2.4.2 Diffusion Pipeline

The Diffusion Pipeline is executed in parallel to the Anatomical Pipeline once the nonuniformity correction stage has completed. If motion correction is enabled, the Diffusion Pipeline will begin by first generating a brain mask using the b = 0 DWI image volumes. This mask is then used by FSL’s eddy (Andersson & Sotiropoulos, 2016) to correct for eddy current distortion and head motion effects in all DWI image volumes. To reduce the possible errors propagated by iterative interpolations, FSL’s eddy estimates both head movement and diffusion eddy current models simultaneously. It also detects slice outliers and optionally removes and replaces the outlier slices using Gaussian process predictions. While motion correction is beneficial when motion or eddy current artifacts are present, it can also degrade image quality if the DWI volumes are poorly registered. In some cases, users may not want to perform these correction steps or may want to perform them using other software prior to running the BrainSuite BIDS App. The default behavior of the Diffusion Pipeline is thus not to apply motion correction, but it is available as an optional step.

The remainder of the Diffusion Pipeline, with the exception of the final two stages, is performed using the standalone BrainSuite Diffusion Pipeline (BDP) executable. BDP first aligns the skull-stripped, nonuniformity-corrected T1w-image generated by the Anatomical Pipeline to the diffusion-weighted image (DWI) data. This is achieved using INVERSION (Bhushan et al., 2012, 2015), a robust image-registration technique that exploits the inverted contrasts between T1- and T2-weighted images. After applying an initial rigid registration to align the DWI and T1w data, INVERSION performs a constrained, non-rigid registration to apply susceptibility-induced geometric image distortion correction without a B_0_ fieldmap. Alternatively, correction using B_0_ fieldmaps can be applied if available. We note that we have not included denoising as part of the Diffusion Pipeline in the BrainSuite BIDS App. A number of programs can be used to preprocess the diffusion data prior to applying the BrainSuite BIDS App, including FSL’s topup (Andersson et al., 2003), as well as our own IPEDcorrect (Bhushan et al., 2013) and joint denoising software (Haldar et al., 2012; Varadarajan & Haldar, 2015). The Diffusion Pipeline provides flexibility by allowing users to perform these DWI corrections using tools outside of the BrainSuite BIDS App prior to running the Diffusion Pipeline. In such cases, the preprocessed data would be considered BIDS Derivatives, thus the BrainSuite BIDS App expects them to be placed separately from input data directory with their locations specified by the user.

BDP next performs tensor fitting and estimation of orientation distribution functions (ODFs) on the coregistered, distortion-corrected data. If the input diffusion MRI data were acquired using a single shell acquisition, then diffusion tensors are fitted and volumetric maps of fractional anisotropy (FA), axial diffusivity (AxD), radial diffusivity (RD), median diffusivity (MD), and apparent diffusion coefficient (ADC) are computed from the fitted tensors. The BDP executable provides multiple methods for estimating ODFs to accomodate a variety of diffusion sampling schemes. The Funk-Radon Transform (FRT; Tuch, 2004) and the Funk-Radon and Cosine Transform (FRACT; Haldar & Leahy, 2013) can both be applied to single-shell data. If FRT is computed, then a volumetric map of generalized fractional anisotropy (GFA) is also calculated from the FRT ODFs. BDP also provides implementations of 3D-SHORE (Özarslan et al., 2013), generalized q-space imaging (GQI) (Yeh et al., 2010), and ERFO (Varadarajan & Haldar, 2017, 2018). 3D-SHORE can estimate ODFs from single-shell and multi-shell data, while GQI can be applied to grid, single-shell, and multi-shell data. ERFO can be applied to single-shell, multi-shell, grid, and arbitrary sampling schemes. The tensor and ODF outputs can be used by several programs, including BrainSuite, to perform tractography and connectivity analysis. We do not currently implement these analyses as part of the BrainSuite BIDS App, though we expect to include them in a future version.

The quantitative diffusion parameter maps (FA, AxD, RD, MD, ADC, GFA) output by BDP can be used in the statistical modeling provided by the group-level analysis stage. In preparation for this, the BrainSuite BIDS App Diffusion Pipeline transforms all computed diffusion parameter maps into atlas space using the mapping produced by the Anatomical Pipeline. Similar to the final smoothing step of the Anatomical Pipeline, the transformed data are smoothed using an isotropic Gaussian kernel with a default size of *σ* = 3 mm. We note that this step is not part of the stand-alone BDP. We also note that the functionality included in the BrainSuite BIDS App Diffusion Pipeline is a subset of features provided by the BDP executable included in the BrainSuite 23a distribution.

#### 2.4.3 Functional Pipeline

The BrainSuite BIDS App will launch the Functional Pipeline once SVReg completes. The Functional Pipeline invokes the stand-alone BrainSuite Functional Pipeline (BFP) executable, which processes raw resting-state or task-based fMRI data to perform motion correction and outlier detection, registers the fMRI data to the corresponding T1w anatomical data, and generates a representation of the fMRI data in grayordinate space in preparation for group-level analysis. BFP is implemented in MATLAB (R2023a) and makes system calls to run BrainSuite, AFNI, and FSL modules to perform these tasks. BFP makes use of a predefined set of surface and volume grayordinates that we have identified on the BCI-DNI atlas that are consistent with the Human Connectome Project (HCP) grayordinate system. These grayordinates were identified by processing the BCI-DNI atlas data with FreeSurfer (Fischl, 2012) and using the spherical maps of surfaces shared by the HCP as described in (A. A. Joshi, Chong, et al., 2018). With these grayordinates established in our BCI-DNI brain atlas space, BFP uses the surface and volume mappings produced by the Anatomical Pipeline to map the subject data to grayordinate space, facilitating group-level analysis.

BFP processes input fMRI data as follows (see Fig. 3). The fMRI data are first deobliqued used AFNI’s 3drefit and reoriented using AFNI’s 3dresample program to be in FSL-compatible, right-posterior-inferior (RPI) orientation (see https://www.fmrib.ox.ac.uk/fsl). A reference image, which is used for motion correction and registration to the T1w data, is then created from the fMRI data using one of three user-selectable methods. The default method, which we have named SimRef, finds an optimal reference image by calculating the structural similarity index (SSIM), a measure of alignment between pairs of images (Hore & Ziou, 2010; Wang et al., 2004), which we compute between every tenth time point and all other time points. The timepoint with the highest mean SSIM, i.e., the highest degree of alignment to the rest of the 4D dataset, is chosen as the reference image. Alternatively, the average of all volumes over the time series can be used as the reference image.

Motion correction is performed using AFNI’s 3dvolreg program to compute a 5-parameter rigid registration of each time point volume in the fMRI data to the reference volume. We perform an initial registration using smoothed images and linear interpolation, followed by a second registration using the original images and Fourier interpolation. BFP then performs motion outlier detection using two methods. The first computes the SSIM between each fMRI time point and the reference image for the original and motion-corrected fMRI datasets. The second uses DVARS from FSL’s fsl_motion_outliers program, which computes the root mean squared difference between time frames N and N+1 in the 4D fMRI data (Power et al., 2012). BFP outputs plots of the SSIM and DVARS values, which the user can review to determine if the amount of motion in the scan is acceptable for the specific study. The outlier volume numbers are provided in a text file that can then be used in GLM or other statistical analysis. The motion-corrected data are then registered to the subject’s T1-weighted image. An initial rigid registration is performed using INVERSION (as described in section 2.4.2). Other cost functions, as well as FSL’s rigid registration tool, are available for optimizing this registration step.

The brain mask produced by the Anatomical Pipeline is transformed from the subject T1w space to the fMRI space. The resampled mask is used to perform skull and scalp removal on the fMRI data. This is followed by spatial smoothing using a Gaussian kernel with a default full-width-half-maximum (FWHM) of 6mm (this can be configured by the user; see App. A.1), which is then followed by grand-mean scaling, temporal bandpass filtering, and detrending. Next, nuisance signal regression, which regresses signals from cerebrospinal fluid, white matter, and motion out of the data, is applied using the FEAT model in FSL (S. M. Smith et al., 2004).

The processed fMRI data are then resampled to 3mm isotropic voxels in the subject’s native T1w space using the registration transform computed between the T1w and the reference image. The data are then transformed to the HCP grayordinate system using the surface-constrained volumetric registration results from the Anatomical Pipeline. Specifically, we use the registration transform inverse to map grayordinate points from the BCI-DNI atlas to the subject T1w space and then resample the fMRI data at those coordinates using linear interpolation. The surface and volume grayordinates are combined to form a vector of size 96,000 (32,000 for each hemisphere + 32,000 for sub-cortex) (Glasser et al., 2013; A. A. Joshi, Chong, et al., 2018). The final output of the individual workflow is a 2D matrix of dimension 96,000 *× time*, where 96,000 is the number of spatial points in grayordinates and *time* is the number of time points in the time series.

Once the fMRI data are transferred into the grayordinate system, we apply a global PDF-based nonlocal means filter (GPDF). GPDF is a data-driven approach for filtering fMRI signals based on the probability density functions (PDFs) of the correlations between time series across spatial points throughout the brain (J. Li et al., 2020). It enables users to perform a global filtering with improved noise reduction effects but without blurring the boundaries between adjacent but distinct functional regions. This enables GPDF to preserve distal and inter-hemispherical correlations.

### 2.5 BrainSuite Dashboard and Quality Control System

We developed a process monitoring and quality control system, termed BrainSuite Dashboard, that integrates with the BrainSuite BIDS App (see Fig. 4). BrainSuite Dashboard is a browser-based system that provides interactive visualization of the intermediate participant-level workflow outputs as they are generated, enabling users to track the state of processing and identify errors as they occur.

**Figure 4:**
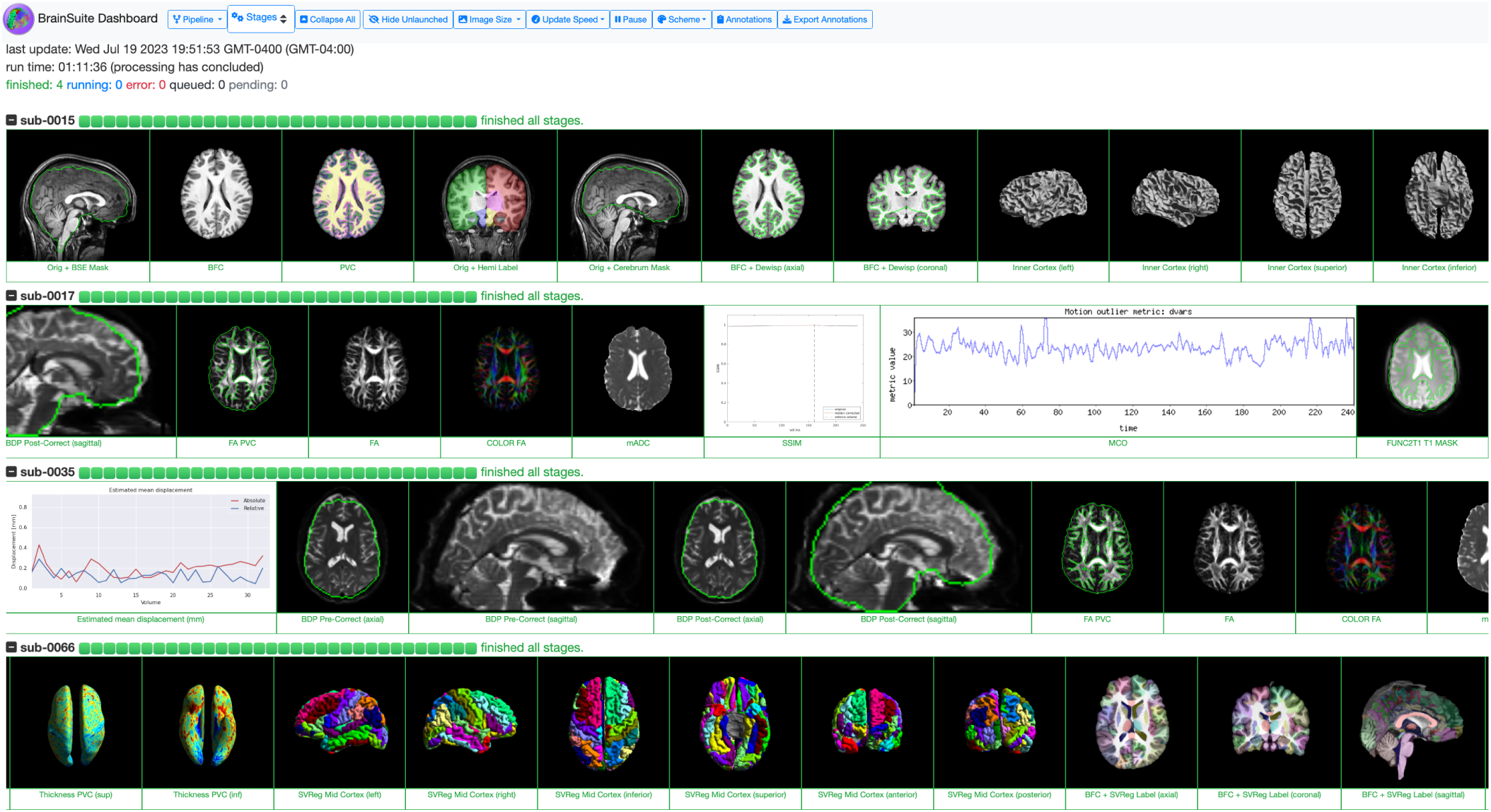
BrainSuite Dashboard. The BrainSuite Dashboard interface provides an interactive view of the participant-level study data as they are processed. Snapshot images of the outputs of key stages are generated by the workflow. The browser view updates as each stage completes, enabling the user to identify processing issues as they occur. The user can select different views that highlight individual pipelines or monitor the overall progress of the data processing for the study. The image row for each participant can be independently expanded and scrolled to facilitate detailed review.

The BrainSuite Dashboard system interfaces with a quality control (QC) component that generates process status updates and output snapshots, which is implemented within the BrainSuite BIDS App. The QC component is invoked as a separate instance that runs concurrently with a set of participant-level workflow instances. Upon initialization, the QC component creates a separate additional output directory, specifiable by the user, that will store files pertaining to the QC system. It then copies to this directory: 1) the BrainSuite Dashboard code, composed of HTML, JavaScript, and CSS files, for interacting with the QC outputs; 2) a JSON file specifying the outputs that are generated and their associated stages in the BrainSuite BIDS App pipelines; 3) a JSON file specifying the dataset description; and 4) a JSON file specifying the subject IDs for the study. The first two components are part of the BrainSuite BIDS App distribution; the second two are generated automatically when the QC system is launched. The QC instance monitors the output directories of the participant-level instances while they are running. The QC instance will periodically (once per second by default) generate a JSON file that represents the current state of processing for all subjects for the entire study. This file captures the current time, the status of the overall processing and the status of processing for each workflow stage for each subject. The status for each subject is encoded as a string of character codes, with each character indicating a stage of processing (e.g., Q for Queued, C for Completed). The QC instance also produces image snapshots of the outputs of workflow stages that we have identified as useful for assessing the quality of BrainSuite outputs. These snapshots are stored in the QC output directory. In most cases, these will be either 2D slices of a 3D image volume or renderings of a surface model. When processing of all subjects has completed, the QC instance terminates.

The outputs of the QC instance can be viewed using BrainSuite Dashboard during and after processing. BrainSuite Dashboard uses JavaScript to provide dynamic content that updates during the participant-level processing. This page must be accessed through a web server due to file system security standards that prevent JavaScript from directly accessing files on the user’s file system, which would prevent BrainSuite Dashboard from reading the JSON files that provide the necessary study, subject, and status information. Many research institutions have web servers attached to a centralized file system that could be used for this purpose. Users can also launch their own web server through various means. The HTTP server built into Python is sufficient for serving the content of BrainSuite Dashboard, and it can be invoked by the user with a single command-line call. This could be launched on a separate computer or from the BrainSuite BIDS App QC instance. Importantly, the web server instance must be run on a system with read access to the output directory and with the ability to serve content through any firewalls between itself and the user. There are a variety of ways to configure such a setup that will be specific to each user’s specific compute environment.

BrainSuite Dashboard loads the study JSON files and, if the processing is ongoing, will read the BrainSuite state JSON file periodically (by default, once per second) to update its content. The user is presented with the overall information about the launched study, including the dataset name, the time started, time of latest update, and information showing how many instances are currently running, queued, completed, or have errors. The user is also presented with a view of the progress for each participant, shown as a progress bar with a node for each stage, and text descriptors of the current state. Each participant row can also be expanded to provide a row of snapshot images depicting the intermediate states of processing. The specific snapshots shown can be selected through dropdown menus that provide selection at the pipeline level (e.g., show all cortical surface extraction outputs) or the individual snapshot level. As new output images are generated by the active participant-level instances, BrainSuite Dashboard updates the image row to include the new snapshots. Users are also provided with an interface for annotating each subject and marking if any subjects should be excluded. The annotation data can be exported as a TSV file, which can be read directly by bstr during group-level analysis. The BrainSuite Dashboard web interface is extensible and can be configured through a JSON file to use different snapshot images and stages, thus enabling additional QC outputs to be added to the system in the future or to apply the BrainSuite Dashboard to different processing pipelines.

### 2.6 Group-level Analysis

Once the participant-level workflow has been completed for all subjects in a study, the user can run the group-level analysis stage of the BrainSuite BIDS App. This requires two additional files: (1) a TSV file containing subject demographic information, such as age and sex, and (2) a JSON file describing the model specification (see Appendix, Fig. 11 for an example). As specified in the BIDS App standard, the first column of the demographics TSV file must be named “participants_id” in the column header. Each row of this column contains the unique subject identifier for that row of data. This subject identifier string should match the exact name of the subject directory that contains the processed data. The other columns are study-specific and provide data for variables collected by the study. Additionally, users can include optional columns that specify whether to exclude individual subjects from the group-level analysis. The model specification file contains user-defined specifications for the type of statistical test, imaging measure, and analysis method used. The types of available tests and metrics are outlined in the following sections.

#### 2.6.1 Group-level Analysis of Anatomical and Diffusion Data

During the group-level analysis stage, outputs from the anatomical and diffusion pipelines are processed using bstr (S. Joshi et al., 2020) to calculate population statistics. We have integrated bstr into the BrainSuite BIDS App by creating an R environment with all the dependencies required for bstr to run. In the BrainSuite BIDS App, bstr is invoked from a Python script using the `rpy2` package. For anatomical data, bstr can perform several analyses, including tensor-based morphometry (TBM) of volumetric data, surface-based cortical thickness analysis, and ROI analysis. For diffusion data, bstr can perform voxelwise analysis of parameter maps (e.g., FA maps), that have been transformed to the volumetric atlas space. Tensor-based morphometry measures the local magnitudes of deformation fields to measure the shrinking and expansions of regions of the brain as it is warped to match another brain. For TBM, this represents the volume change that each voxel in the atlas undergoes when it is deformed to the corresponding region in the subject image. Cortical thickness comparisons map the thickness measures from the subject cortical mesh to the vertices of the atlas cortical mesh, enabling vertex-wise comparison across the study population. ROI analysis can be performed on structural data. It compares regional statistics of mean grey matter (GM) volume, mean white matter (WM) volume, or mean GM thickness within anatomical ROIs defined by the labeling produced by the Anatomical Pipeline.

The statistical tests provided by bstr include Pearson correlation, general linear model, ANOVA, t-test, and permutation tests. The population statistical tests are performed at each voxel for TBM and diffusion parameter map comparisons, and at each vertex for surface-based thickness analysis. For ROI-based analysis, these tests are performed on grey matter and white matter volume measures computed within ROIs or on cortical thickness measures averaged over cortical ROIs. The available group-level analysis outputs and available statistical tests are summarized in Table 1.

For surface-based analyses, the outputs generated by bstr include separate right and left mid-cortical surface maps in the atlas space that store unadjusted and adjusted t-values and log p-values thresholded by level of significance (p < 0.05) at each vertex. For volumetric-based analyses, adjusted and unadjusted t-value and log p-value volumetric overlays and corresponding lookup table (LUT) files are produced. The LUT files, when loaded into the BrainSuite GUI, follow the same color scheme as the surface files. Annotated bi-directional colorbars are also rendered as PDF files, which can be integrated directly into publications.

For volumetric and ROI analyses, bstr also provides automated report generation to visualize statistical results using R-shiny and R Markdown. Examples of bstr reports are shown in Figures 6, 7, and 9. The volumetric analysis report displays a table that lists peak coordinates of clusters (contiguous significant voxels). It also displays renderings of the clusters for both the unadjusted and the adjusted versions of the p-values and the t-statistics, which are overlaid on corresponding slices from the T1-w MRI of the atlas to provide neuroanatomical context. The report provides an interface to sort these results by the size of the cluster or the peak values of the t-statistics. The ROI analysis report shows the demographic spreadsheet, automatic bar plots for ANOVA and regressions, and scatter plots for correlation analyses. Additionally, bstr exports an R Markdown report that contains reproducible R commands in both the RMD file and the HTML document (Xie, 2015). This enables complete reproducibility of statistical results and only requires that the R Markdown file is packaged along with the BIDS formatted data needed for bstr.

#### 2.6.2 Running group-level analysis on outputs from the Functional Pipeline

The stand-alone BrainSuite Functional Pipeline (BFP) provides novel statistical methods to perform group-level analysis using BrainSync, an orthogonal transform that performs temporal alignment of time-series at homologous locations across subjects allowing direct comparison of scans (A. A. Joshi, Chong, et al., 2018). Correlations between time-series on spatially homologous points are measured across subjects or between a subject and atlas. FMRI atlases are created within the group-analysis workflow using BrainSync Alignment (BSA), a method we developed that jointly synchronizes fMRI data across the time-series data of multiple subjects (Akrami et al., 2019). BFP’s group-level analysis is called from a Python script and reads grayordinate BFP outputs, a demographic spreadsheet in TSV format, and a configuration file in JSON format. Results are generated in grayordinate space, with labeled cortical representations generated on BrainSuite surface files and noncortical results written out as 3D NIfTI files.

The fMRI statistical analysis pipelines include novel pair-wise regression and group differences tools (see Tab. 1). These include tools for group-wise BrainSync that synchronize fMRI signal from multiple subjects (Akrami et al., 2019) to generate group fMRI atlas, pair-wise regression, and group differences (A. A. Joshi et al., 2021). These pipelines facilitate population studies of fMRI for resting-state and task-based paradigms. In group studies, especially in the case of spectrum disorders, distances to a single atlas do not fully reflect the differences between subjects that may lie on a multi-dimensional spectrum. Our pair-wise approach measures the distances between pairs of subjects and uses these distances as the test statistics rather than comparing subjects to the group mean or to a single reference point or template. This approach has been shown to be more sensitive in localizing the group effects (A. A. Joshi et al., 2021).

##### BrainSync

BrainSync is an orthogonal transformation that enables comparison of resting fMRI (rfMRI) time series across subjects. This method exploits similarity in correlation structure across subjects to perform an orthogonal rotation of time-series data between two or more subjects to induce a high correlation between time series at homologous locations. The fMRI data at each vertex are normalized to zero mean and unit norm. After normalization of the time series at each vertex, each time series can be represented as a point on a hypersphere. Under the assumption of similar spatial correlation patterns across subjects, it was previously shown (A. A. Joshi, Chong, et al., 2018) that between every pair of subjects, there exists an orthogonal transform that approximately synchronizes their time-series data. Because similar connectivity patterns are observed across subjects, a single orthogonal transformation can be performed to minimize the geodesic distance between homologous points across subjects. The geodesic distance between homologous points is observed to compute a measure of similarity between subjects. The orthogonal transformation that performs the synchronization is unique, invertible, efficient to compute, and preserves the connectivity structure of the original data for all subjects. Similar to image registration, where we spatially align the anatomical brain images, this synchronization of brain signals across a population or within subject across sessions facilitates connectivity studies of rfMRI data. In contrast with existing fMRI analysis methods, this transform does not involve dimensionality reduction and preserves the rich functional connectivity information in rfMRI scans.

##### Atlas-Based Statistical Analysis

For each subject, the atlas-based BFP statistical analysis synchronizes fMRI data of the atlas and the subject, and then computes measures of similarity between the atlas and the subject’s time-series data at each point in the grayordinate system. BrainSync Alignment (BSA), which jointly synchronizes fMRI data across the time-series data of a pool of representative subjects (Akrami et al., 2019), is used to create a custom fMRI atlas. The reference atlas can be created from the entire study population or from a subset of it, such as subjects within the control group. Alternatively, a representative individual dataset can be identified that has the lowest pair-wise distances between all pairs of subjects, and those data can be used as the reference. The atlas-based statistical analysis performs Pearson correlations and controls for covariates using linear regression. The correlation coefficient and unadjusted and adjusted p-values (FDR) are reported.

##### Pair-wise Statistical Analysis

The pair-wise BFP statistical analysis performs statistical tests for associations and group differences using pair-wise distances between random pairs of subjects’ fMRI data as well as the pair-wise differences between the corresponding test variable or group indicators. The pair-wise method for statistical analysis may be particularly useful in studies where it is difficult to determine which subjects or group should be assigned as the representative data. Pearson correlation is performed to test for associations, and t-tests are performed to test for group differences. P-values are corrected for multiple comparisons using FDR or max-T permutations. Both unadjusted and adjusted p-values are reported for pair-wise statistical testing. The outputs generated by these analyses include surface and volumetric files in the grayordinate space containing R-values and unadjusted and adjusted p-values (FDR and max-T; Nichols & Holmes, 2002).

### 2.7 Documentation and User Support

Detailed end-user documentation is provided at https://brainsuite.org/BIDS/, where users can find descriptions of the pipelines and workflows in the BrainSuite BIDS App, installation and usage instructions, walk-through demos for running participant-level processing, and the complete set of instructions for reproducing the results presented in this manuscript.

User support for the BrainSuite BIDS App is provided in three ways: 1) through emails to our email address; 2) through our BrainSuite support forums, which includes a new forum dedicated to the BrainSuite BIDS App; and 3) through the issue tracker associated with the BrainSuite BIDS App GitHub page.

## 3 Results

We applied the BrainSuite BIDS App to data from the Amsterdam Open MRI Collection’s (AOMIC) Population Imaging of Psychology (PIOP) cohort (Snoek et al., 2021) to demonstrate its functionality and capabilities. The AOMIC PIOP datasets include imaging and behavioral subject data collected from healthy participants, all of whom were university students. As detailed below, we first processed these data using the participant-level workflows. We then evaluated the outputs using the BrainSuite Dashboard system, tuned program settings based on this evaluation, and reprocessed the participant-level data with the new settings. We then performed five types of group-level analysis on the outputs of the participant-level processing: a) surface-based thickness analysis; b) tensor-based morphometry; c) ROI analysis; d) functional connectivity analysis; and e) fractional anisotropy analysis. For the anatomical analyses (a-c), we examined the effects of Raven’s Advanced Progressive Matrices (RAPM) scores (Raven & Court, 1938; Spielberger et al., 1968; Van der Ploeg, 1982), a proxy measure for intelligence, on brain structure. For the fMRI analysis (d), we examined the effects of RAPM scores on functional connectivity.

For the diffusion MRI analysis (e), we performed a comparison of sex differences in fractional anisotropy values. We selected these particular analyses because these topics are well-studied and the outputs of our software would be expected to be concordant with the published literature. Furthermore, to differentiate between the statistical effects of the RAPM scores according to sex, all analyses (except the analysis of sex differences in FA) were performed within separate male and female cohorts.

### 3.1 Data

Data were pooled from the two AOMIC PIOP (Snoek et al., 2021) datasets: PIOP1 (N = 216 ; age 22.18 ± 1.80 years, 18.25 to 26.25; 120 F / 89 M / 7 unspecified), which was collected in 2016, and PIOP2, which was collected in 2017 (N = 226; age 21.96 ± 1.79 years, 18.25 to 25.75; 129 F / 96 M / 1 unspecified). All images in PIOP1 and PIOP2 were acquired on the same Philips 3T scanner. However, there was a system upgrade between these projects, with the PIOP1 dataset being acquired using the “Achieva” version and the PIOP2 being acquired using the “Achieva dStream” version. We retrieved the AOMIC PIOP datasets from the OpenNeuro database (Snoek et al., 2020a, 2020b). We excluded 23 subjects that were missing demographic data (N=10) or imaging data (N=13), resulting in a total number of 419 subjects for our analyses (N=419, age 22.05 ± 1.79 years; 240 F / 179 M). We used the subject demographic variables for age, sex, and Raven’s Advanced Progressive Matrices (RAPM) scores collected for the AOMIC study. RAPM scores are designed to measure general and fluid intelligence using nonverbal abstract reasoning problems (Raven & Court, 1938; Spielberger et al., 1968; Van der Ploeg, 1982). A higher RAPM test score indicates higher accuracy in the participants’ Raven’s matrices responses, indicating greater fluid intelligence.

### 3.2 Ethics Statement

All data used in this study were open-access data retrieved from the OpenNeuro archive (Markiewicz et al., 2021) and used under a CC0 - Creative Commons public domain license. Specifically, we made use of the AOMIC-PIOP1 (Snoek et al., 2020a) and AOMIC-PIOP2 (Snoek et al., 2020b) datasets, which were deidentified and made publicly available by Snoek et al. (2021). As described in the reference paper for these data (Snoek et al., 2021), the acquiring study was approved by the local ethics committee at the University of Amsterdam (EC number: 2010-BC-1345) and conducted with the informed consent of all participants.

### 3.3 Participant-level Processing

All participant MRI data were first processed using BrainSuite BIDS App’s participant-level processing pipeline using the default parameters and the default BCI-DNI Brain Atlas, with the exception of two processing options: T1w rescaling during skull stripping in the Anatomical Pipeline and applying motion correction during the Diffusion Pipeline. The T1w rescaling was performed during skull stripping by the Brain Surface Extractor (BSE) module, which normalizes the intensity values of the T1w data to the range [0,32000] using the formula *I_norm_* = *max*(0, (32000 *× I*)*/max*(*I*)), where *I* is the set of image intensity values. We evaluated the quality of the initial participant-level outputs using the BrainSuite BIDS App Dashboard QC system. The Dashboard showed poor cerebrum labeling in most cases, leading to poor cortical boundary extraction. Based on this, we modified two parameter settings for cerebrum labeling (initializing with centroids and linear convergence value) and repeated the image processing and QC evaluation for all subjects. Inspection of the QC results indicated that the processing was suitable for 406 of the 419 subjects. We identified 13 subjects whose data were were not processed successfully. Three of these cases were corrected by selecting subject-specific skull-stripping parameters (anisotropic diffusion constant and edge detection constant). Five cases required adjustments to the cerebrum labeling parameters (initializing with centroids and cost function type). For the remaining five subjects, we changed the cost functions used for rigid registration during the Functional Pipeline. We then reran the BrainSuite BIDS App on these 13 subjects, resulting in complete participant-level outputs for all N=419 subjects.

### 3.4 Group-level Analysis

We performed an ANOVA examining the main effects of Raven’s score on anatomical structures and functional connectivity on the AOMIC-PIOP dataset. We also investigated differences in fractional anisotropy (FA) values between the two sexes. In all tests, handedness was not considered as a covariate because it did not differ significantly across sex. All tests were corrected for multiple comparisons using FDR, with the exception of ROI analysis, which considered only one ROI under the family of ROI-wise statistical tests.

#### 3.4.1 Surface-Based Analysis

We performed an initial surface-based analysis (SBA) of cortical thickness, which was then used to determine the appropriate tests to be performed for the ROI, TBM, and FC analyses, described in the following sections. We first performed an ANOVA on the pooled dataset (N = 419; age 22.05 ± 1.79 years; 240 F / 179 M) that investigated the effect of RAPM scores (24.47 ± 4.853) on cortical thickness measures while controlling for age, sex, and scanner type. The results showed a larger effect size and greater surviving effects on both hemispheres in the female group (N = 240; age 22.07 ± 1.74 years, 18.25 to 26.25) when compared to the male group (N = 178; age 22.04 ± 1.86 years, 18.25 to 26.25).

Two new ANOVA analyses examining the effect of RAPM scores on cortical thickness measures while controlling for age and scanner type were next performed separately on the female and male groups. In the female group, we observed significantly lower cortical thickness proportional to increasing Raven’s score; these effects were seen diffusely throughout the brain (see Fig. 5). The largest and most significant clusters were observed bilaterally in the superior and middle frontal gyri and orbitofrtonal cortex. In addition, large clusters were observed in the left inferior frontal gyrus, left visual cortex, right paracentral lobule and right cingulate cortex, which were correlated with Raven’s score after vertex-wise FDR correction. The male cohort did not exhibit any significant clusters after FDR correction.

**Figure 5:**
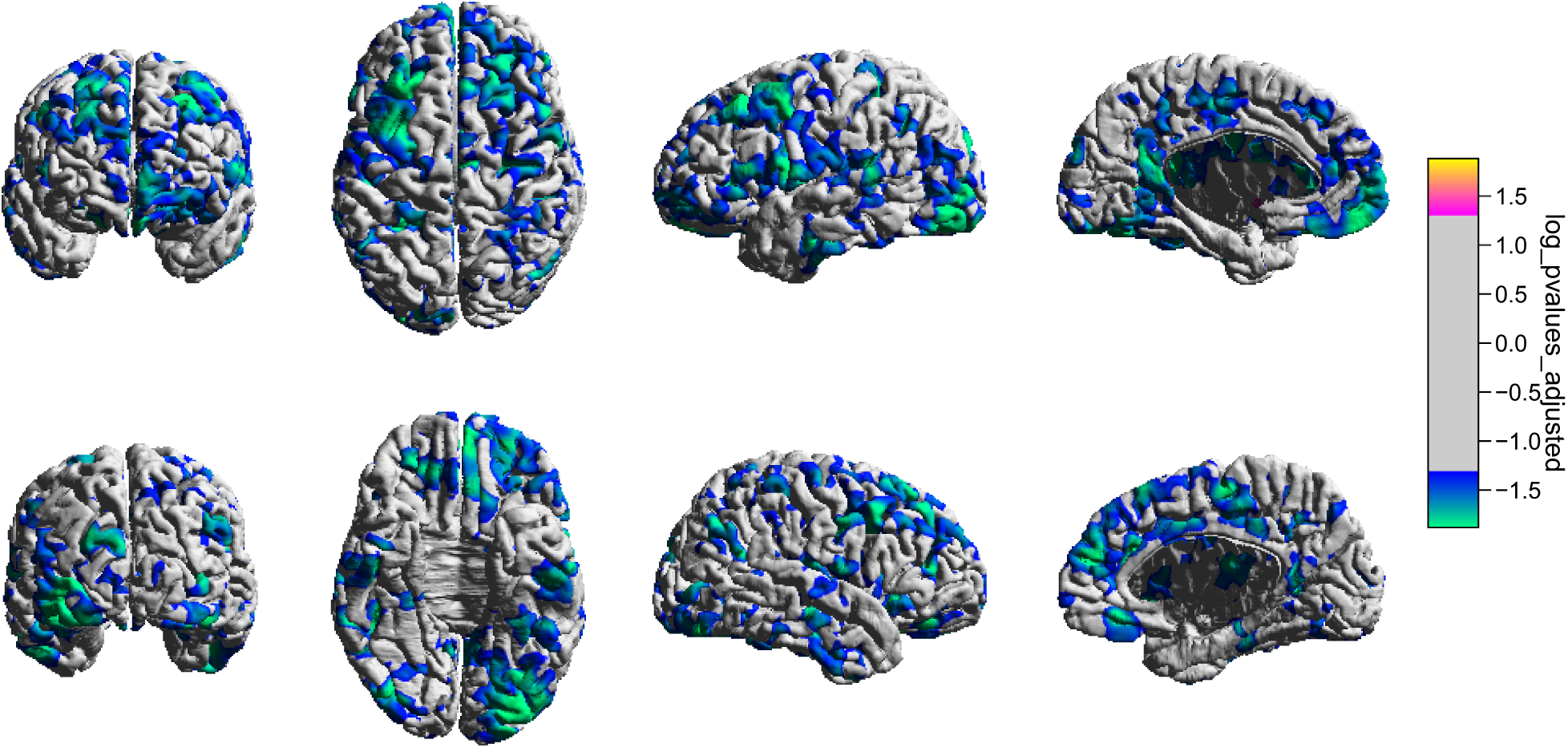
Surface-based analysis of the effects of RAPM scores on cortical thickness in females in the AOMIC-PIOP dataset. Shown are the outputs of a surface-based analysis performed using bstr, which examined the effects of RAPM scores on cortical thickness in females in the AOMIC-PIOP dataset. The color bar on the right indicates the adjusted log transformed p-values after FDR correction: blue hues in the color bar denote significantly lower cortical thickness proportional to increasing RAPM scores, whereas pink hues indicate the opposite. The colors shown on the cortex in these views indicate regions where female participants had lower cortical thickness proportional to higher RAPM scores.

#### 3.4.2 Tensor-Based Morphometry Analysis

Based on the results of the surface-based analysis, we focused our subsequent analyses, with the exception of the FA analysis, on the female cohort. Using bstr, we performed a post-hoc ANOVA investigating the main effect of RAPM scores on TBM in females. The results revealed no statistical significance after FDR correction. We show the uncorrected results in Fig. 6, which includes a truncated version of the bstr report, to illustrate the functionality of the BrainSuite BIDS App and its components. The largest cluster comprised 37,516 voxels with an average t-value of 3.628 and was located in the white matter interior to the posterior portion of the left superior temporal gyrus, indicating a possible increase in volume in this region with increasing RAPM scores. The second largest cluster (13,702 voxels; average t-value = 2.989) was located in the right orbitofrontal cortex, again suggesting an increase in volume in this area with increasing RAPM scores. None of these voxels survived significance testing after FDR correction.

**Figure 6:**
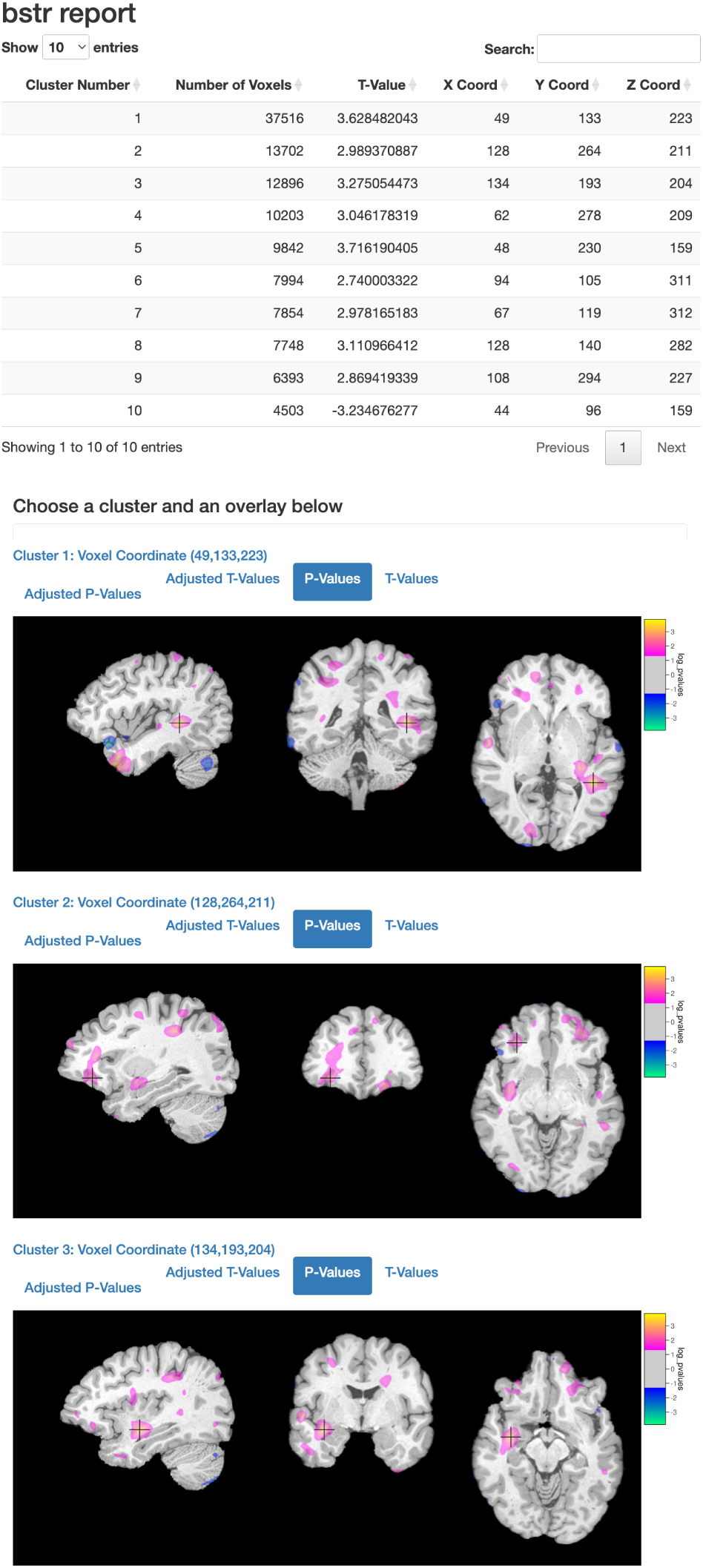
Tensor-based morphometry analysis of the effects of RAPM score on local volume in females in the AOMIC-PIOP dataset. Shown is a bstr report detailing the results of a TBM analysis of the effects of the RAPM score on local volume in the female cohort. The first ten largest clusters are listed in a table at the top of the report. The blue hues indicate reduction in volume with increasing RAPM scores and pink hues indicate increase in volume with increasing RAPM scores. The test produced no surviving significant voxels after FDR correction, thus, we show the unadjusted log-transformed p-value maps. The largest cluster of 37,516 voxels with an average t-value of 3.628 is located in the white matter interior to the posterior portion of the left superior temporal gyrus.

#### 3.4.3 ROI Analysis

Surface-based analysis results (Fig. 5) showed significantly lower cortical thickness in multiple areas. We selected one of these areas, the left pars opercularis, for ROI analysis. Using bstr, we conducted an ANOVA to study the effects of RAPM scores on the average grey matter thickness values for this ROI. Figure 7 shows the results of the ANOVA and a scatter plot for this study, captured directly from the report generated by bstr. Coinciding with our finding in SBA, these results indicate that grey matter thickness values decreased with increasing RAPM score values (F-statistic = 10.5; p-value = 0.0014).

**Figure 7:**
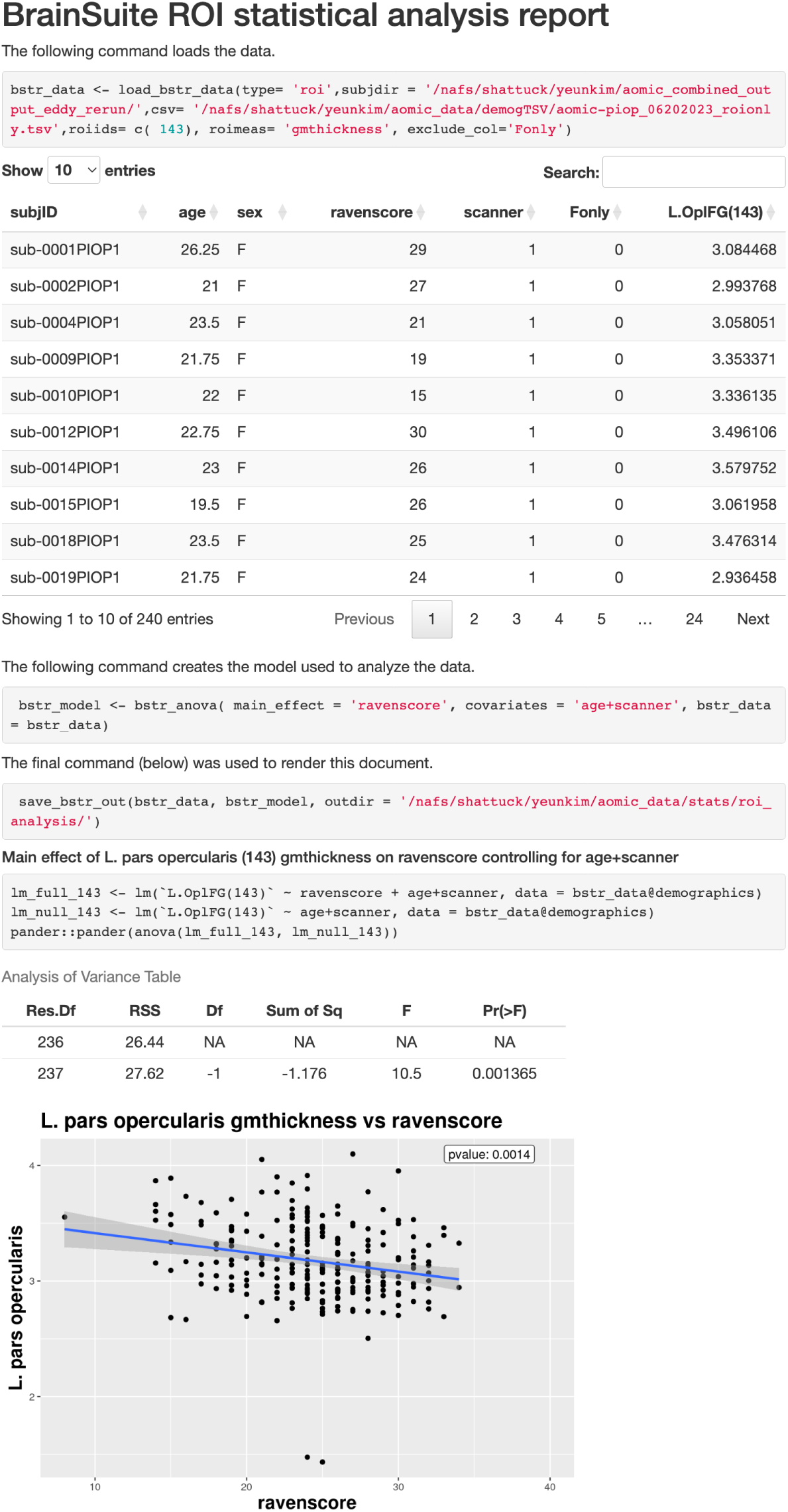
ROI analysis of the association of grey matter thickness in the left pars operularis with RAPM score in females in the AOMIC-PIOP dataset. Shown is the entire bstr report, which begins with a table of the demographic data and the exact commands used in the statistical test. This is followed by a table of the summary statistics and a scatter plot that shows significantly lower GM thickness with increasing RAPM score values.

#### 3.4.4 Functional Connectivity Analysis

Using BFP’s group analysis tools, we performed an atlas-based linear regression test on the effects of RAPM score on functional connectivity. The atlas was generated using data from the 13 subjects with the highest scores (range: 32-35) on the RAPM test. We hypothesized greater functional differences in brain connectivity patterns in relation to increasing differences in RAPM scores. We therefore used FC data from subjects with high RAPM scores as the reference for the linear relationship. Thus for the specific test used here, positive R-values, which correspond to a higher geodesic distance to the atlas, indicate that a higher RAPM score was associated with lower similarity to the atlas. Similarly, negative R-values, which correspond to a lower geodesic distance to the atlas, indicate that a higher RAPM score was associated with higher similarity to the atlas.

The cortical functional connectivity results are shown in Fig. 8. No significant clusters survived after FDR correction. In the uncorrected R-value results, we see that a relationship between functional connectivity and RAPM score was found diffusely throughout the brain, especially along the bilateral medial frontal gyri, right dorsolateral prefrontal cortex, anterior cingulate cortices, right temporal pole, left middle temporal gyri, bilateral inferior temporal gyri, right precuneus, and occipital areas. The blue areas indicate regions where a higher RAPM score was associated with higher similarity to the atlas.

**Figure 8:**
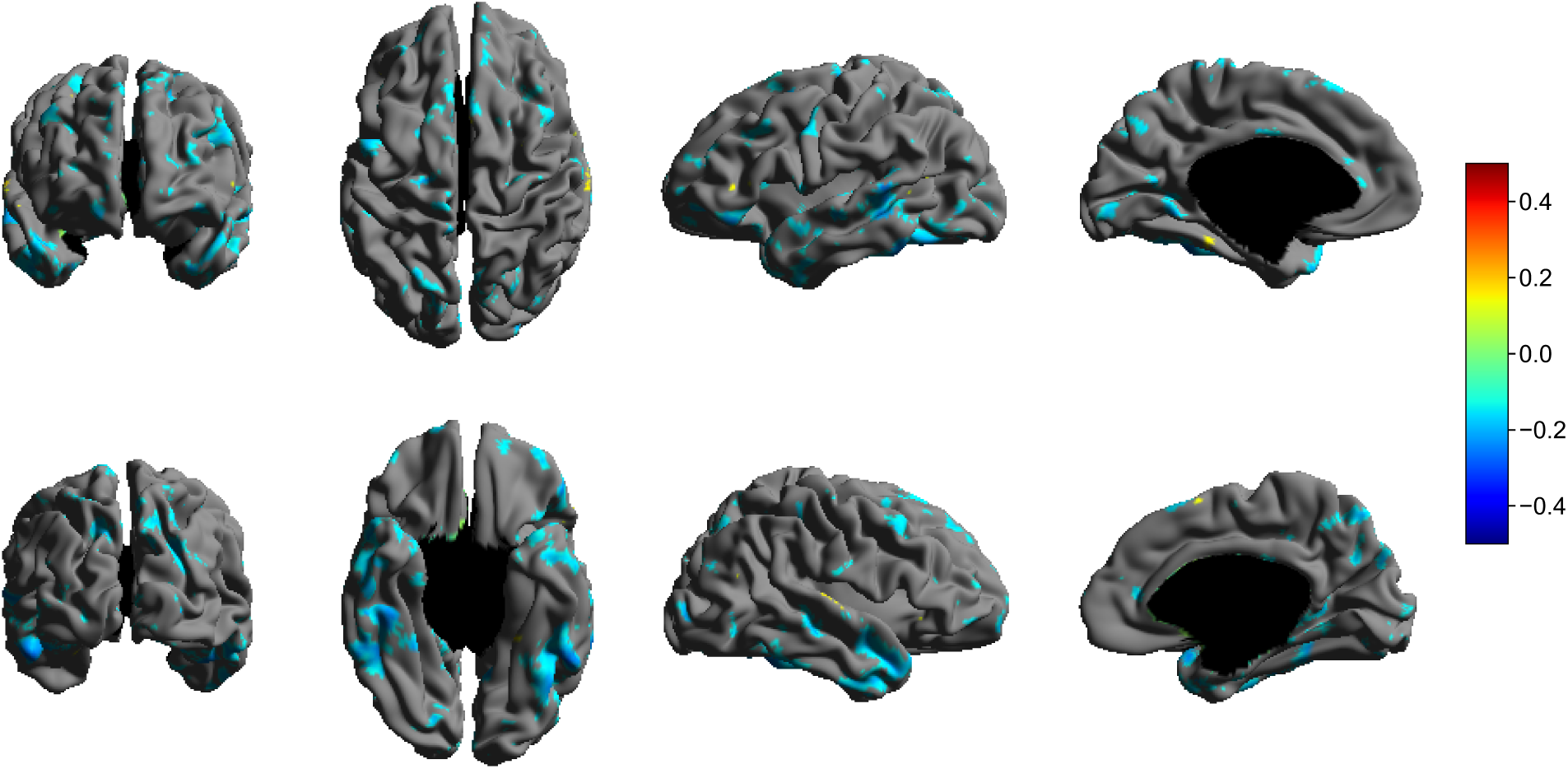
Functional connectivity analysis of the effect of RAPM score on cortical functional connectivity in the female group. Shown are unadjusted R-values from an ANOVA analysis. The color bar on the right indicates a symmetric range of +0.5 (red) to −0.5 (blue) R-values. Positive R-values indicate that a higher RAPM score was associated with lower similarity to the atlas, while negative R-values indicate that a higher RAPM score was associated with higher similarity to the atlas. The atlas was created using FC data from the 13 subjects with the highest RAPM scores. Areas associated with Raven’s scores included the bilateral medial frontal gyri, right dorsolateral prefrontal cortex, anterior cingulate cortices, right temporal pole, left middle temporal gyri, bilateral inferior temporal gyri, right precuneus, and occipital areas.

#### 3.4.5 Fractional Anisotropy Analysis

We next analyzed voxel-wise differences in FA values between males and females (N = 419; age 22.05 ± 1.79 years; 240 F / 179 M) using a t-test. The bstr analysis report for this study is shown in Fig. 9. The largest cluster consisted of 106,660 voxels and was located in the right globus pallidus (t-value = −8.863) and extended into the right thalamus, the right external capsule, the right putamen, and the right caudate nucleus. The next largest cluster was located in the left occipital lobe, with 22,125 voxels and an average t-value of −5.932. In both of these clusters, males were observed to have higher FA values relative to females. However, in the third largest cluster, located in the left paracentral lobule (21,355 voxels and average t-value= 5.512), females were observed to have higher FA values.

**Figure 9:**
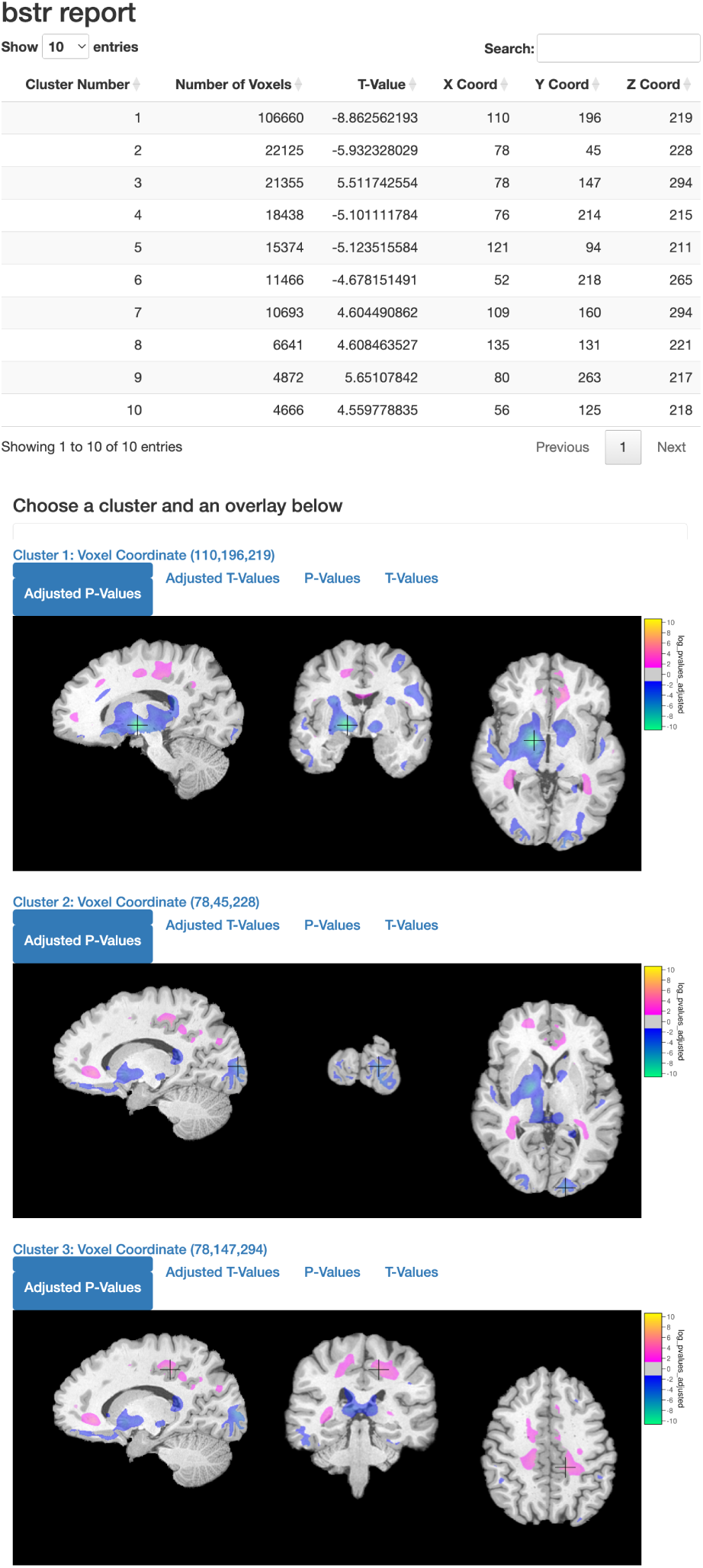
Fractional anisotropy analysis examining sex differences in FA values using a t-test on the AOMIC-PIOP dataset. Shown is the bstr report presenting the analysis of sex differences in fractional anisotropy conducted using a using a t-test on the AOMIC-PIOP dataset. The first column includes a table of ten significant clusters. Images of the adjusted log p-value maps overlaid on the BCI-DNI brain atlas follow the table. Blue hues indicate lower FA values and pink hues indicate higher FA values in females when compared to males. The images display the center of the identified cluster, indicated by the black cross-hair, in all three cardinal planes. The largest cluster, comprising 106,660 voxels with a peak occurring at the right globus pallidus (t-value = −8.863) and extending into the right thalamus, the right external capsule, the right putamen, and the right caudate nucleus.

### 3.5 Computer Resource Allocation and Benchmarks

The data processing described in the preceding sections was performed on a multi-user high performance computing (HPC) cluster operated by the UCLA Brain Mapping Center. The participant-level workflows were distributed on the cluster and executed in parallel using SLURM (Yoo et al., 2003) to perform job scheduling and maintenance. Each Apptainer instance was allocated 64 GB of memory and 7 processing cores, which were assigned by SLURM from a pool of compute nodes with either dual 12-core 2.6GHz Intel Xeon Gold 6126 or dual 18-core 2.3GHz Intel Xeon Gold 6140 CPUs. Execution of the full workflow, including the Anatomical Pipeline, the Diffusion Pipeline, the Functional Pipeline, and the Quality Control component, required approximately 125 minutes total time for each subject. Approximately 22 workflows were run concurrently, enabling all participant-level workflows to be completed in approximately 40 hours of real time. Processing of the data from four participants used to generate Fig. 4 was performed in parallel on a single Intel i9-13900K Debian 12 workstation with 128GB of RAM and required less than 70 minutes. When run serially, processing required less than 55 minutes per participant dataset on the same computer.

All group analyses were also performed on the UCLA Brain Mapping Center cluster, with resource allocation that varied depending on the type of analysis. For SBA, 60 GB of memory and 10 cores were allocated, allowing the entire analysis to complete in less than 4 minutes. For TBM and FA analyses, 150 GB of memory and 10 cores were allocated, and approximately 9 minutes were required for each analysis. For ROI and FC analyses, 10 GB of memory and 6 cores were allocated. The ROI analysis and bstr report generation required approximately 1 minute and FC analysis required approximately 24 minutes.

## 4 Interpretation of AOMIC Results

The series of studies we conducted during the evaluation of our BrainSuite BIDS App on the AOMIC dataset on cortical thickness, ROI, TBM, and FC focused on the effects of RAPM scores in females. These studies yielded findings similar to those in the existing literature. The statistical tests for these models were performed separately within the male and female groups to disassociate effects due to sex. The statistical analyses in the female group showed bigger effect sizes and survived multiple testing on both hemispheres. Although our TBM and FC analyses did not produce statistically significant results, we included the uncorrected findings to demonstrate the functionality of our software and to show the reporting functionality of bstr. We also performed a separate study that included the entire dataset and examined sex differences on FA measures.

### 4.1 Surface-based and ROI analysis

The results of our cerebral cortical thickness analysis and our ROI analysis of the pars opercularis both corroborate the findings reported by Schnack et al. (2015) and Selemon (2013). In those studies, grey matter thickness measures were shown to decrease with RAPM scores, a phenomenon the authors suggested could be caused by cortical pruning. Moreover, a study by Shaw et al. (2006) showed a delayed increase in cortical thickness in children with higher intelligence scores when compared to other youth with varying levels of intelligence, possibly indicating a longer period of cognitive circuit development.

### 4.2 Tensor-based morphometry (not significant)

Our TBM analysis studying the relationship between grey matter volume and RAPM scores in the female cohort showed small, statistically insignificant changes in the cortical regions. There are some regions for which the results are in line with the findings from the surface-based analysis (e.g., bilateral orbitofrontal cortices), but some clusters (e.g., left temporal pole) found in the TBM analysis are not observed in SBA. It appears that some of the effects seen in this TBM analysis are within the white matter regions. This is reasonable, because white matter volume has been associated with fluid intelligence (Turken et al., 2008). Additionally, our results showed higher volume in the regions of the salience network, including the medial frontal lobe, regions of the temporal lobes, and the sensorimotor cortex, which aligns with the results presented by Yuan et al. (2012). That study used a similar number of subjects as our study and did obtain a surviving significance; in contrast to our method, they investigated only grey matter and used Monte Carlo simulation for multiple comparisons correction. Lastly, we note that the number of voxels compared in the TBM study was far greater than the number of vertices compared in the surface-based analysis, which led to a larger number of statistical tests.

### 4.3 Functional Connectivity (not significant)

Although the evidence seen in the functional connectivity analysis was not statistically robust, the directionality of the R-values, especially in the temporal lobe, did correspond to findings in previous studies. The temporal lobe has been shown to be associated with inhibitory control and attention through its WM connections to the frontal lobe (Ramezanpour & Fallah, 2022). Significant associations have been observed between Flanker test scores (a proxy measure of cognition) and temporal lobe cortical thickness (Sarabin et al., 2023), and a recent study suggested that white matter BOLD activity in the temporal lobe mediates age-related cognitive changes (M. Li et al., 2024). In agreement with these findings, we observed differences (though not statistically significant) in functional connectivity in the temporal lobe in subjects associated with RAPM scores in our study, suggesting that RAPM tests may involve similar cognitive functioning as Flanker tests.

### 4.4 Fractional Anisotropy

Previous studies investigating sex differences in FA measures have reported significant effects in multiple anatomical regions. Several studies have reported mixed findings for differences in FA in terms of the direction (higher or lower) and the specificity of the regions (Chou et al., 2011; Inano et al., 2011; Menzler et al., 2011). Our findings showed higher FA in males in the thalamus, hypothalamus, globus pallidus, and putamen in agreement with Menzler et al. (2011) and den Braber et al. (2013), and lower FA in the anterior cingulate cortex, the fronto-occipital fasciculus and white matter under the parahippocampal gyrus similar to Abe et al. (2010) and Chou et al. (2011). The differences in FA measures may be related to variations in the proportion of grey and white matter across the sexes (Allen et al., 2003; Goldstein et al., 2001). Additionally, myelination rate has been found to be enhanced by estrogen hormones in young rat brains (Prayer et al., 1997). Thus, it is possible that a similar mechanism could be present in human brains and modulate the FA measures.

## 5 Discussion

BrainSuite is one of many freely available tools for analyzing neuroimaging data. Several existing packages have also been integrated into BIDS Apps, in some cases by their own development teams and in some cases by interested third parties. Additionally, there have been efforts to develop BIDS Apps that create new pipelines by integrating software tools developed by multiple groups. We briefly describe a few major packages that share similarities in features and functions with the BrainSuite BIDS App.

One important category of BIDS Apps are those developed by the NeuroImaging PREProcessing toolS (NiPreps) project, which aims to design minimal preprocessing pipelines for neuroimaging data. The FMRIPrep BIDS App (Esteban et al., 2019) contains standardized pipelines for processing raw fMRI data using tools from various software packages, including AFNI (Cox, 1996), FSL (S. M. Smith et al., 2004), FreeSurfer (Fischl, 2012), and ANTs (Avants et al., 2011). FMRIPrep performs structural preprocessing, coregistration of fMRI and T1w data, fMRI reference image estimation, fMRI image unwarping (i.e. head motion, slice time, susceptibility distortion correction), and resampling to native T1w and standard spaces using methods from various packages. Similarly, QSIprep consists of workflows for processing raw DWI images using methods from software tools from several developers. Although QSIprep’s pipeline has dMRI-specific processing methods, it shares some components of the processing steps from fMRIPrep (e.g., FreeSurfer-based structural data processing and ANTs SyN-based fieldmapless distortion correction (Esteban et al., 2019)) and also employs AFNI, FSL, FreeSurfer and ANTs. QSIprep is suitable for use with various diffusion sampling schemes, and it performs denoising, distortion correction, and head motion correction. It also performs registration and normalization to T1w space if a T1w MRI is provided. In the final stages of their pipeline, QSIprep offers many options for diffusion model fitting and connectivity matrix computation using tools from Dipy (Garyfallidis et al., 2014), MRtrix (Tournier et al., 2019), and DSI Studio (Yeh & Tseng, 2011). Both fMRIPrep and QSIprep generate quality control reports that users can use to identify outliers in their datasets.

A comprehensive quality assessment can be performed using the MRIQC BIDS App (Esteban et al., 2017, 2022), also from the NiPreps group. The MRIQC BIDS App provides a series of tools for assessing image quality at the individual and group level. MRIQC produces image quality metrics (IQMs) using a combination of FSL, ANTs, and AFNI. It generates a report for each subject at the participant-level, as well as a group report including all subjects. The group report presents boxplots and stripplots of the IQMs, enabling the identification of subject data that may be of poor quality. We note that these tools are more comprehensive than the quality control provided in our BrainSuite BIDS App.

While fMRIPrep, QSIprep, and MRIQC BIDS Apps provide unique methods and features for fMRI and dMRI data processing, respectively, and quality assessment, the BrainSuite BIDS App offers the capability to process T1w, dMRI, and fMRI data and to perform group analysis, all within the same container. By including participant- and group-level workflows, the BrainSuite BIDS App allows users to perform end-to-end structural, diffusion, and functional MRI analysis from one program.

A number of BIDS Apps have also been developed that are focused on existing individual software packages. The FreeSurfer recon-all BIDS App (Fischl, 2012; K. J. Gorgolewski et al., 2018) is a containerized version of the FreeSurfer cortical surface reconstruction pipeline. It takes T1-weighted images as input and generates registered cortical surfaces, cortical and subcortical regional parcellations, and measures including cortical thickness. If multi-session data are provided, it will run the FreeSurfer longitudinal pipeline. The FreeSurfer BIDS App provides two group-level workflows, which (1) create subject-specific templates; and (2) generate data tables of all cortical and subcortical segmentation statistics and image quality statistics. While the FreeSurfer BIDS App does not include FreeSurfer’s group-level statistical analysis tools, modeling can be performed outside of the BIDS App using mri_glmfit or its corresponding front-end GUI QDEC (Query, Design, Estimate, Contrast) application. A separately developed BIDS App (Liem & Gorgolewski, 2017) implements an interface to FreeSurfer’s TRACULA (Yendiki et al., 2011), which provides tools for extracting white matter pathways by processing dMRI data from cross-sectional and longitudinal datasets. TRACULA can perform FreeSurfer’s cortical surface reconstruction pipeline, but has limited flexibility compared to the FreeSurfer BIDS App. For its group-level analysis, TRACULA BIDS App outputs motion and tract statistics for all subjects for users to run statistical modeling and inference externally.

Though not part of the BIDS App framework, the FreeSurfer toolset also provides tools for analyzing image and segmentation quality, including the FreeSurfer 5.3 Quality Assessment (QA) Tools (https://surfer.nmr.mgh.harvard.edu/fswiki/QATools) and their replacement qatools-python (https://github.com/Deep-MI/qatools-python). These tools are compatible with FreeSurfer outputs and provide a range of functions, which include the generation of a limited set of thumbnail images of processed images at specific stages and computation of quality assessment metrics (QAMs), e.g., signal-to-noise ratio (SNR), segmentation outliers, and topological holes or errors.

The MRtrix3 Connectome BIDS App (R. E. Smith, 2018; R. E. Smith & Connelly, 2019), based on MRtrix3 (Tournier et al., 2019), provides a fully automated pipeline that estimates structural connectomes and prepares them for group comparison. It requires one or more series of DWI images and one T1w-MRI. DWI data are processed using a series of operations, including anatomically constrained tractography (R. E. Smith et al., 2012) and application of SIFT2 (R. E. Smith et al., 2015) to produce streamlines.

Streamlines are assigned to grey matter parcellations created from the T1w-MRI using either FreeSurfer or atlas-based registration. MRtrix3 Connectome BIDS App’s group-level analysis normalizes connection densities across subjects so that connectome edges can be compared. It also produces an FA-based population template and a group mean connectome.

The Statistical Parametric Mapping (SPM) BIDS App (Flandin et al., 2018) provides an instance of SPM12 (Ashburner, 2012), which provides a broad range of functionality for analyzing many types of data, including T1w-MRI, fMRI, positron emission tomography (PET), single photon emission computed tomography (SPECT), electroencephalography (EEG), and magnetoencephalography (MEG) data. It also offers the ability to coregister these datasets with T1-weighted images for anatomical interpretation. SPM BIDS App is structured to take and execute a user-written MATLAB script for the group-level analysis and an optional MATLAB script for preprocessing. These scripts might typically make use of the spm_jobman functionality, which enables the specification of a series of SPM processes to be linked together into a pipeline. This offers a great deal of flexibility for performing different types of analysis, but also requires substantial scripting. This is in contrast to the FreeSurfer, MRTrix3, and BrainSuite BIDS Apps, which each contain pre-programmed pipelines that can be invoked with a single command-line call.

We developed the BrainSuite BIDS App to provide access to the comprehensive functionality of the major processing capabilities available in BrainSuite. Specifically, the integration of the three BrainSuite pipelines into a single BIDS App provides a greatly simplified process for applying these methods to study data maintained in BIDS format. While these pipelines are available through sets of scripts, command-line programs, and, in the case of the BrainSuite Anatomical Pipeline, an interactive GUI, the BrainSuite BIDS App enables rapid deployment for large-scale analysis and ensures consistent versioning of the software and the environment in which it resides. Furthermore, we have designed the BrainSuite BIDS App to interoperate with the command-line and GUI versions of BrainSuite, enabling users to perform customized processing or perform manual actions, such as mask editing, if necessary. We have also introduced new functionality in the BrainSuite BIDS App that is not currently included in the main BrainSuite distribution.

One of the improvements we made in developing the BrainSuite BIDS App was the direct integration of the BrainSuite Statistics in R toolbox into our BIDS analysis workflow. This creates fully automated analysis streams for structural and diffusion data, from processing of individual datasets to complete statistical analysis reports. We note that most of the BIDS Apps described above, with the exception of SPM, prepare data for analysis during the group-level stage, but the final statistical modeling is performed in separate tools outside of the BIDS App. Importantly, our BrainSuite framework captures the full set of parameters used across an entire study in a set of JSON files. The statistical analysis performed is also recorded in the files output by bstr. These archival mechanisms enhance interpretation and reproducibility by ensuring that the processing and analysis are well-documented. In addition, the integration of the BrainSuite Functional Pipeline and the BrainSync group analysis tools provides a novel mechanism for synchronizing functional signals across individuals in a study.

The BrainSuite BIDS App also introduces the BrainSuite Dashboard quality control system, which provides an interactive interface to monitor and review intermediate stages of subject-level data processing as they are completed. This is in contrast to the methods of MRIQC, which runs a series of steps to produce a quality report prior to analysis, or FreeSurfer, which provides tools outside of the BIDS or BIDS App frameworks that generate post hoc reports after the recon-all process has completed. An advantage of the MRIQC and FreeSurfer approaches is that they compute quantitative metrics of quality. While our current approach relies upon visual inspection, we note that we have previously explored computing quality assurance measures for use in an earlier version of our QC system (Wong & Shattuck, 2018). We plan to continue that work to develop quality assurance modules for BrainSuite that will identify potential processing errors and abnormalities. These would then be flagged for review in the BrainSuite Dashboard. We also note that the BrainSuite Dashboard is designed to be easily modified. The pipelines, stages, and images shown to the user are all configurable through JSON files. This enables new processing modules to be added to the system with minimal effort. In a separate project, for example, we have reconfigured this system for use in a preclinical imaging pipeline. This framework could also be adapted to work with the outputs of other BIDS Apps.

As we demonstrated through the processing of the open-access AOMIC-PIOP data, the integration of our BrainSuite pipelines and functionality into a single container in the form of a BIDS App provides a convenient and powerful mechanism for researchers to process and analyze large-scale neuroimaging studies. By following the BIDS App model, the BrainSuite BIDS App is designed to promote reproducibility and interoperability. It maintains a consistent virtual environment on any supported platform, requiring users to install only the Docker or Apptainer software and the BrainSuite BIDS App image. Additionally, by using Nipype as the underlying workflow framework, BrainSuite BIDS App benefits from Nipype’s built-in command archiving, logging, and error detection protocols. The Nipype logs supplement BrainSuite’s own logs and command archival, thereby enhancing data provenance and error assessment. These features improve usability by alleviating the need for configuring computer environments and by providing the user with additional information regarding the status of processing. As our BrainSuite software continues to evolve, we plan to integrate and distribute these updates into the BrainSuite BIDS App to further facilitate reproducible neuroimaging processing and analyses.

## Data and Code Availability

The full set of data used in this manuscript are available for download from OpenNeuro (AOMIC-PIOP1 v2.0.0 [https://openneuro.org/datasets/ds002785/versions/2.0.0] and AOMIC-PIOP2 v2.0.0 [https://openneuro.org/datasets/ds002790/versions/2.0.0]). All additional files necessary to reproduce the analyses performed in this manuscript (demographic files formatted for use with BrainSuite BIDS App, JSON specification files, and intensity rescaling software) are available from https://github.com/BrainSuite/BrainSuiteBIDSAppPaperData. A detailed description of how to rerun the analyses is available on our website https://brainsuite.org/BIDS/paper. We have also packaged a subset of data from 4 participants and made it available from https://github.com/BrainSuite/BrainSuiteBIDSAppSampleData for demonstration purposes. These data were used to produce Fig. 4. An interactive demo of the BrainSuite Dashboard interface using this dataset is also provided on our GitHub site (https://brainsuite.github.io/DashboardDemo/).

The BrainSuite BIDS App code is open source and freely available from https://github.com/BIDS-Apps/BrainSuite. A pre-built Docker image containing the BrainSuite BIDS App is freely available from https://hub.docker.com/r/bids/brainsuite. Additionally, a pre-built Apptainer-based BrainSuite BIDS App is available for download at http://brainsuite.org/data/sif/bids_brainsuite_v23a.simg. The BrainSuite BIDS App makes use of multiple free and open-source packages, which are detailed in the BrainSuite BIDS App GitHub repository.

## Author Contributions

**Yeun Kim**: Data curation, Software, Formal analysis, Investigation, Methodology, Validation, Visualization, Writing - original draft, Writing - review & editing; **Anand Joshi**: Software, Methodology, Writing - original draft, Writing - review & editing; **Soyoung Choi**: Software, Methodology, Writing - original draft, Writing - review & editing; **Shantanu Joshi**: Software, Methodology, Writing - original draft, Writing - review & editing; **Chitresh Bhushan**: Software, Methodology, Writing - review & editing; **Divya Varadarajan**: Software, Methodology, Writing - review & editing; **Justin Haldar**: Software, Methodology, Writing - review & editing; **Richard Leahy**: Software, Funding acquisition, Methodology, Supervision, Writing - review & editing; **David Shattuck**: Conceptualization, Software, Funding acquisition, Methodology, Project administration, Resources, Supervision, Validation, Visualization, Writing - original draft, Writing - review & editing.

## A Customization

### A.1 Participant-level Workflow

Users can run the participant-level workflow with custom parameters by including a preprocessing configuration file (preproc_config.json; see Fig. 10). This optional JSON file defines objects named Anatomical, Diffusion, PostProc, and Functional that can be used to specify parameters for each of those components. For parameters whose options are Boolean values, 1 (True) and 0 (False) are used, e.g., 1 enables an option and 0 disables it.

#### A.1.1 Global Settings

◦ `cacheFolder`: Specifies the location of the cache directory for Nipype. The default location for the cache directory is the output directory.

#### A.1.2 Anatomical Pipeline Settings

Parameters for the BrainSuite BIDS App Anatomical Pipeline are set in the Anatomical object of the JSON config file. There are no required user-defined settings for the Anatomical Pipeline.

Skull-stripping (bse)

◦ `skipBSE`: If enabled (1), skull stripping (BSE) is skipped and a custom mask is used (e.g., if manual edits to a brain mask were necessary). The custom mask must be copied to the output directory prior to running the Anatomical Pipeline. Options: {0,1}. Default: 0 (disabled).
◦ `autoParameters`: Enables automated selection of optimal parameters for skull-stripping/Brain Surface Extractor (BSE). Options: {0,1}. Default: 1 (enabled).
◦ `prescale`: Prescales image to uint16 data type for skull-stripping/Brain Surface Extractor (BSE). Options: {0,1}. Default: 0 (disabled).
◦ `diffusionIterations`: Specifies the number of times the anisotropic diffusion filter is applied to the image during BSE. This field will be ignored if autoParameters is 1. Typical range: [0*…* 6]. Default: 3.
◦ `diffusionConstant`: Specifies the anisotropic diffusion constant, which controls the height of the edges that are retained during anisotropic diffusion filtering. This field will be ignored if autoParameters is 1. Typical range: [5*…* 35]. Default: 25.

**Figure 10:**
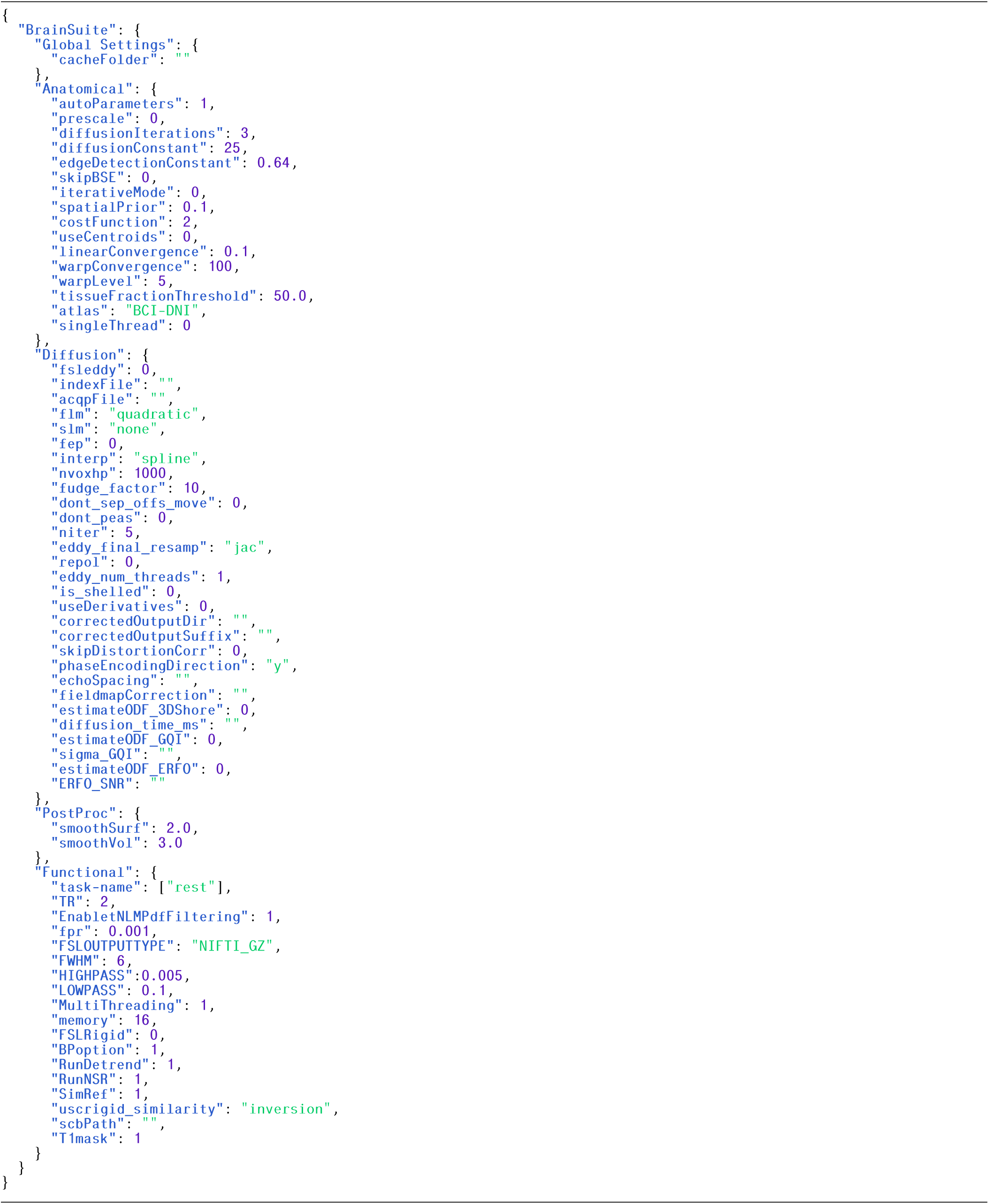
Example JSON file defining the preprocessing specification for BrainSuite BIDS App.
◦ `edgeDetectionConstant`: specifies the edge detection constant, *σ*. During the edge detection step in BSE, *σ* influences how wide an edge must be to be identified. Typical range: (0.5-1.0). Default: 0.64

Bias Field Correction (bfc)

◦ `iterativeMode`: If iterative mode is enabled, the bias field correction (BFC) program will run multiple passes using a range of settings to correct for severe artifacts. Options: {0,1}. Default: 0.

Tissue Classification (pvc)

◦ `spatialPrior`: Controls the weighting of the spatial prior used during tissue classification stage. Reducing this value can be useful if an image has low signal-to-noise. Default: 0.1.

Cerebrum Labeling (cerebro)

◦ `costFunction`: The cost function used by AIR’s alignlinear during the initial linear registration to the atlas for cerebrum labeling. Possible choices are: standard deviation of the ratio image (0), least squares (1), and least squares with intensity rescaling (2). Options: {0,1,2}. Default: 2.
◦ `useCentroids`: If enabled, the cerebrum labeling program will initialize the registration process using the centroids of the subject image and the atlas image. Options: {0,1}. Default: 0.
◦ `linearConvergence`: Sets the threshold used by AIR’s alignlinear during cerebrum labeling to determine if the linear registration has converged. Default: 0.1.
◦ `warpConvergence`: Sets the threshold used by AIR’s align_warp during cerebrum labeling to determine if the nonlinear registration has converged. Default: 100.
◦ `warpLevel`: Sets the degree of the polynomial model used for the transformation used by AIR’s align_warp during cerebrum labeling. Typical Range: (2-8). Default: 5

White Matter Mask Generation (cortex)

◦ `tissueFractionThreshold`: Minimum percentage of white matter in a voxel needed for it to be included in the mask, in decimal form (e.g., 50% white matter) during initial cortical mask generation. Range: (0-100). Default: 50.0

Registration and Labeling (svreg)

◦ `atlas`: Specifies the atlas used for registration and labeling (see 2.4.1 for details on the atlases). Options: BSA, BCI-DNI, USCBrain.^1^ Default: BCI-DNI.
◦ `singleThread`: If enabled, SVReg runs in single-threaded mode by disabling multithreading in MATLAB’s parpool. This can be helpful if errors related to MATLAB parpool occur on compute nodes. Options: {0,1}. Default: 0

#### A.1.3 User-Defined Configurations for the Diffusion Pipeline

Parameters for the Diffusion Pipeline are set in the Diffusion object of the JSON config file. There are no required user-defined inputs for the Diffusion Pipeline if FSL’s eddy (eddy current and motion correction) is disabled. If FSL’s eddy is enabled, users must specify acqpFile and indexFile fields.

The following parameters in the Diffusion Pipeline pertain to FSL’s eddy (Andersson & Sotiropoulos, 2016). For more information regarding these arguments, with the exception of fsleddy and eddy_num_threads, please visit https://fsl.fmrib.ox.ac.uk/fsl/fslwiki/eddy/UsersGuide.

◦ `fsleddy` : If enabled, FSL eddy (Andersson & Sotiropoulos, 2016) will perform eddy current and motion correction Options: {0,1}. Default: 0.
◦ `acqpFile` : Path to a user-defined text file that contains diffusion data acquisition parameter information (e.g., phase encoding direction, blip-up/blip-down, and pixel/bandwidth). This parameter is required when fsleddy (FSL eddy) is enabled.
◦ `indexFile` : Path to user-defined text file that indicates which volumes in the diffusion data follow which acquisition parameters written in the text file in the acqpFile field. This parameter is required when fsleddy (FSL eddy) is enabled.
◦ `flm` : Sets the model for estimating the eddy current-induced field. This parameter is used when fsleddy is enabled. Options: {linear,quadratic,cubic}. Default: quadratic.
◦ `slm` : Sets the model for estimating the eddy currents caused by diffusion gradients. This parameter is used when fsleddy is enabled. Options: {none,linear,quadratic}. Default: none.
◦ `fep` : If enabled, FSL eddy identifies and fills empty planes. This parameter is used when fsleddy is enabled. Options: {0,1}. Default: 0.
◦ `interp` : Sets the model for interpolation during resampling. This parameter is used when fsleddy is enabled. Options: {spline,trilinear}. Default: spline.
◦ `nvoxhp` : Number of voxels to use during the hyperparameter estimation of the Gaussian Process during the prediction phase. This parameter is used when fsleddy is enabled. Default: 1000.
◦ `fudge_factor` : Sets the level of Q-space smoothing during the prediction phase. This parameter is used when fsleddy is enabled. Default: 10.
◦ `dont_sep_offs_move` : If enabled, FSL eddy will not attempt to distinguish the constant component of the eddy current field and subject movement and will only fit one model (the constant component of the eddy current field). This parameter is used when fsleddy is enabled. Options: {0,1}. Default: 0.
◦ `dont_peas` : If enabled, FSL eddy will not estimate the movement between the b = 0 volume and the first diffusion data volume. This parameter is used when fsleddy is enabled. Options: {0,1}. Default: 0.
◦ `niter` : Specifies the number of iterations. This parameter is used when fsleddy is enabled. Default: 5.
◦ `eddy_final_resamp` : Sets the method (Jacobian modulation or least-squares reconstruction) for final resampling. This parameter is used when fsleddy is enabled. Options: {jac,lsr}. Default: jac.
◦ `repol` : If enabled, replaces outlier slices using predictions from Gaussian Process. This parameter is used when fsleddy is enabled. Options: {0,1} Default: 0.
◦ `eddy_num_threads` : Sets the number of threads used for FSL eddy. Default: 1.
◦ `is_shelled` : If enabled, FSL eddy automatically assumes that the diffusion data are shelled. This parameter is used when fsleddy is enabled. Options: {0,1}. Default: 0.

The following set of parameters pertain to the stand-alone BrainSuite Diffusion Pipeline (BDP; Varadarajan et al., 2020).

◦ `skipDistortionCorr`: If enabled, BDP skips susceptibility-induced geometric image distortion correction completely and performs only a rigid registration of the diffusion and T1-weighted images. This can be useful when the input diffusion images do not have any susceptibility-induced geometric image distortion or they have already been corrected for the distortion. Options: {0,1}. Default: 0.
◦ `phaseEncodingDirection`: Sets the phase encoding direction of the DWI data, which is the dominant direction of distortion in the images. This information is used to constrain the susceptibility-induced geometric image distortion correction along the specified direction. Directions are represented by any one of x, x-, y, y-, z or z-. x direction increases towards the right side of the subject, while x-increases towards the left side of the subject. Similarly, y and y-are along the anterior-posterior direction of the subject, and z and z-are along the inferior-superior direction. When this field is not specified, BDP uses y as the default phase-encoding direction. Options: {x,x-,y,y-,z,z-}. Default: y.
◦ `echoSpacing`: Sets the echo spacing in units of seconds, which is used during fieldmap-based distortion correction. (Example: For an echo spacing of 0.36 ms, use echo-spacing = 0.00036). This value is required when using fieldmapCorrection.
◦ `fieldmapCorrection`: Use an acquired fieldmap for distortion correction. The parameter specifies the path to the field map file to use.
◦ `diffusion_time_ms`: Sets the diffusion time parameter (in milliseconds). This parameter is required for estimating ERFO, 3D-SHORE and GQI ODFs.
◦ `estimateODF_3DShore`: If enabled, estimates ODFs using the 3D-SHORE (Özarslan et al., 2013) basis representation. Options: {0,1}. Default: 0.
◦ `estimateODF_GQI`: Estimates ODFs using the GQI method (Yeh et al., 2010). Options: {0,1}. Default: 0.
◦ `sigma_GQI`: Sets the GQI adjustable factor, required for calculating diffusion sampling length. Typical range: [1*…* 1.3]. Default: 1.25.
◦ `estimateODF_ERFO`: If enabled, estimates ODFs using the ERFO method. (Varadarajan & Haldar, 2018). Options: {0,1}. Default: 0.
◦ `ERFO_SNR`: Sets the SNR of the acquired data, required for estimating ERFO ODFs. Default: 35.

Users are welcome to preprocess their DWI images prior to running the Diffusion Pipeline using programs external to the BrainSuite BIDS App. The preprocessed outputs, which will be used as inputs to the Diffusion Pipeline, can then be specified in the following fields.

◦ `useDerivatives` : If enabled, the Diffusion Pipeline will use diffusion data derivatives defined by the fields correctedOutputDir and correctedOutputSuffix. This allows users to preprocess their diffusion data outside of the BrainSuite BIDS App and use them as inputs for the Diffusion Pipeline. Options: {0,1}. Default: 0.
◦ `correctedOutputDir` : The path to the directory that contains the preprocessed diffusion data. The subject IDs must be the same as those in the input BIDS data input directory and must adhere to the following folder organization: “derivatives_directory/sub-subjectID/dwi/”. This parameter is required when useDerivatives is enabled.
◦ `correctedOutputSuffix` : The file suffix of the preprocessed diffusion data. Since the preprocessing methods can vary, you can defined the file suffix in this field. The suffix contains all the characters after the subject ID but does not include the file extension. For example, for file “sub-01_dwi.topup.corr.nii.gz”, the file suffix is “_dwi.topup.corr”. The file suffix must be the same for the diffusion image, the b-value text file, and the b-vector text file. This parameter is required when useDerivatives is enabled.

#### A.1.4 Smoothing Parameters

Parameters for smoothing of surface and volume are set in the PostProc object of the JSON config file.

◦ `smoothSurf`: Specifies the kernel size (in mm) used for smoothing the surface output data from the Anatomical Pipeline. Typical range: (2-5). Default: 2.
◦ `smoothVol`: Specifies the kernel size (in mm) used for smoothing the volumetric output data from the Anatomical Pipeline and the Diffusion Pipeline. Typical range: (2-6). Default: 3.

#### A.1.5 User-Defined Configurations for Functional Pipeline

Parameters for the Functional Pipeline are set in the Functional object of the JSON config file. These are passed to the BrainSuite Functional Pipeline (BFP) executable. BFP requires one field, TR, to be specified by the user. This can be set as an argument on the command line when the BrainSuite BIDS App instance is started or set in the preproc-config JSON file. If TR is specified on the command line, then the Functional section of the JSON file is optional. If TR is not defined when the BrainSuite Functional Pipeline is called, then the default (2) will be used. Additional optional fields can be used to control different aspects of the processing.

◦ `task-name`: The names of the tasks to be processed in list form using square brackets, e.g., [restingstate, emomatching]. The task names should correspond to the task names specified in the input fMRI filenames after the task-delimiter. For example, in the fMRI file sub-0001_task-restingstate_bold.nii.gz, the task name would be ‘restingstate’.
◦ `TR`: Repetition time (in seconds) of the fMRI data.
◦ `EnabletNLMPdfFiltering`: If enabled, BFP will apply tNLMPdf (GPDF) filtering (J. Li et al., 2020). This step can take up to 30 minutes per scan. Options: {0,1}. Default: 1.
◦ `fpr`: False positive rate (significance level). This parameter is used for global non-local means filtering (GPDF) for fMRI denoising. Default: 0.001
◦ `FSLOUTPUTTYPE`: Specifies the format that FSL uses to save its outputs.^2^ Default: NIFTI_GZ
◦ `FWHM`: Full-width-half-maximum value, in mm, used for spatial smoothing. Default: 6
◦ `HIGHPASS`: Value for the high-pass cutoff frequency, in Hz, used for bandpass filtering. Default: 0.005
◦ `LOWPASS`: Value for the low-pass cutoff frequency, in Hz, used for bandpass filtering. Default: 0.1
◦ `MultiThreading`: If enabled, uses parallel processing for transforming fMRI data onto the grayordinate system and GPDF non local means filtering. If disabled, parallel processing is not used. {0,1}. Default: 1
◦ `memory`: Specifies the amount of RAM (in gigabytes) available for running for transforming fMRI data onto grayordinate system and GPDF non local means filtering. Default: 16
◦ `FSLRigid`: If enabled, BFP uses FSL’s rigid registration (FLIRT) during processing. If not enabled, BFP uses BrainSuite’s BDP affine registration. FLIRT is run with 6 degrees of freedom, trilinear interpolation, and the correlation ratio cost function (Jenkinson et al., 2002). BrainSuite’s affine registration tool is set to run with 6 degrees of freedom, linear interpolation, and the INVERSION cost function, which is optimized to align the inverted contrasts of T1W and fMRI images (Bhushan et al., 2015). Options: {0,1}. Default: 0
◦ `SimRef`: Specifies the type of reference volume used for coregistration and motion correction. If enabled, SimRef will be used, which calculates the pair-wise structural similarity index (SSIM) between every tenth time point and all other time points (Hore & Ziou, 2010; Wang et al., 2004). The time point with the highest mean SSIM is chosen as the reference image. If not enabled, all volumes are averaged together to create a mean image. {0,1}. Default: 1
◦ `RunDetrend`: Enables detrending. Options: {0,1}. Default: 1
◦ `RunNSR`: Enables nuisance signal regression. Options: {0,1}. Default: 1
◦ `uscrigid_similarity`: Specifies the cost function(s) used by BFP during USC rigid registration. Available methods are INVERSION (inversion), INVERSION followed by normalized mutual-information based refinement (Bhushan et al., 2015) (bdp), mutual information (mi), correlation ratio (cr), and squared difference (sd). Options: {bdp, inversion, mi, cr, sd}. Default: inversion
◦ `T1mask`: If enabled, BFP uses the T1w mask to threshold fMRI data, which may be useful for data with high signal dropout. Options: {0,1}. Default: 1
◦ `BPoption`: If enabled, BFP applies 3dBandpass (updated function with quadratic detrending). If not enabled, BFP applies 3dFourier and linear detrending. Details are found in the AFNI documentation^3^. Options: {0,1}. Default: 1

### A.2 Group-level Analysis

A JSON file specifying a model is required for group level-statistical tests. Example JSON files for configuring group-level analysis / models are shown in Fig. 11. For parameters whose options are Boolean values, 1 (True) and 0 (False) are used.

#### A.2.1 Group level analysis for anatomical and diffusion data

◦ `tsv_fname`: Specifies the TSV file containing demographic data and/or clinical variables that will be used for group analysis
◦ `measure`: Specifies the imaging measure of interest. Available options are: cortical thickness measures from surface files (sba); tensor-based morphometry using the Jacobian determinant of the deformation map from subject to atlas (tbm); ROI-based analysis using scalar summary statistics (roi); DTI parametric maps (dba). Options: {sba, tbm, roi, dba}
◦ `test`: Specifies the model. Available options are: ANOVA (anova); correlations test (corr); t-test (ttest). Options: {anova, corr, ttest}
◦ `main_effect`: Specifies the main predictor variable for ANOVA. Used only if test=anova. This value must match a corresponding column header in the TSV file that is specified in the tsv_fname field.
◦ `covariates`: Specifies the covariates for ANOVA. Used only if test=anova. Values must match column headers from the TSV file in the tsv_fname field. Must be in list form and can contain multiple elements.
◦ `corr_var`: Specifies the variable for correlation test. Used only if test=corr.
◦ `paired`: Specifies the t-test type. If enabled (1), then paired t-test will be performed. If disabled (0), then unpaired t-test will be performed. Used only if test=ttest.
◦ `group_var`: Specifies the group variable for paired t-test. Used only if test=ttest and paired=1.
◦ `smooth`: Specifies the smoothing level used for sba, tbm, and dba. The smoothing levels must match the levels used during preprocessing. For example, if surface smoothing levels were set to 2.0mm for participant-level processing, then sba’s smoothing level should be set to 2.
◦ `mult_comp`: Specifies the method for multiple comparison correction. Used only if measure is sba, tbm, or dba. Available options are: FDR (fdr); max-T permutation test (perm). Options: {fdr, perm}
◦ `niter`: Specifies the number of iterations for the permutation method. Only used if mult_comp=perm.
◦ `pvalue`: Specifies the method for computing p-values. Available options are: parametric method, which is the classical p-value method (parametric); permutation method, which is the Freedman-Lane method (Freedman & Lane, 1983) (perm). Options: {parametric, perm}
◦ `roiid`: BrainSuite ROI ID number for ROI analysis. Only used if measure= roi. Must be in list format. Multiple IDs can be listed. For example, to study the left (641) and right thalamus (640): [640,641].
◦ `hemi`: (For measure: sba) Hemisphere selection. Options: {left, right, both}
◦ `maskfile`: (For measure: tbm, dba) If mask file is specified, then only the regions within the mask will be considered for statistical analysis. The mask file must be in the same space as the atlas that was used to register the subjects during SVReg.
◦ `atlas`: Atlas file. Default is the atlas that was used to run SVReg.
◦ `roimeas`: (For measure: roi) ROI measure of interest. Available options are: average GM thickness in cortical regions (gmthickness); average GM volume (gmvolume); average wm volume (wmvolume). Options: {gmthickness, gmvolume, wmvolume}
◦ `dbameas`: (For measure: dba) Diffusion measure of interest. Options: {FA, MD, axial, radial, mADC, FRT_GFA}
◦ `exclude_col`: User can specify column header name in the TSV file (the same TSV file in the tsv_fname field). This column must exist in the TSV file for each subject/row, in which 0 indicates no exclusion and 1 indicates exclusion
◦ `out_dir`: Output directory location where statistical analysis results will be stored. This directory will be created if it does not exist.

#### A.2.2 Group-level analysis for functional data

◦ `tsv_fname`: Name of the TSV file containing demographic data and/or clinical variables that will be used for group-level analysis.
◦ `file_ext`: Input file suffix, which is the BFP output in grayordinate space, e.g., ’-rest_bold.32k.GOrd.filt.mat’.
◦ `lentime`: Number of timepoints in the fMRI data.
◦ `matchT`: If some subjects have less than timepoints (’lentime’), enabling this field will add zero values to match number of timepoints. Options: {0, 1}
◦ `stat_test`: Model to be run. Options:{atlas-linear, atlas-group, pairwise-linear}
◦ `pw_pairs`: (For stat_test: pairwise-linear) Number of random pairs to measure.
◦ `pw_fdr`: (For stat_test: pairwise-linear) Multiple comparisons correction method. If enabled (1), FDR correction will be used. If disabled (0), max-T permutations will be used.
◦ `pw_perm`: (For stat_test: pairwise-linear and pw_fdr: False) Number of permutations used for max-T permutation method.
◦ `outname`: File prefixes for statistical output files.
◦ `sig_alpha`: P-value significance level (alpha).
◦ `smooth_iter`: Level of smoothing applied on brain surface outputs
◦ `save_surfaces`: If enabled (1), save surface files. If disabled (0), do not save surface files. Options: {0, 1}
◦ `save_figures`: If enabled (1), to save PNG snapshots of the surface files. If disabled (0), does not save. Options: {0, 1}
◦ `atlas_groupsync`: If enabled, an atlas is generated by first performing group alignment of fMRI data and then averaging over the entire group. If disabled, a reference atlas is created by identifying one representative subject. Options: {0, 1}
◦ `atlas_fname`: File name of user-defined atlas. Variable should be called atlas_data. Leave empty if no user-defined atlas should be used
◦ `test_all`: If enabled (1), subjects used for atlas generation are included during hypothesis testing. If disabled (0), subjects used for atlas creation are excluded from testing your hypothesis. Options: {0, 1}
◦ `colvar_main`: For linear regression or group testing, the main effect of study.
◦ `colvar_reg1`: For group comparisons. assign all rows with zero values if running linear regression. Control up to 2 variables by linearly regressing out the effect. If you only have less than 2 variable you would like to regression out, you can create and assign a dummy column(s) with zero values for all rows.
◦ `colvar_reg2`: Same as colvar_reg1.
◦ `colvar_exclude`: User can specify column header name in the TSV file (the same TSV file in the tsv_fname field). This column must exist in the TSV file for each subject/row, in which 0 indicates no exclusion and 1 indicates exclusion.
◦ `colvar_atlas`: User can specify column header name in the TSV file (the same TSV file in the tsv_fname field). This column must exist in the TSV file for each subject/row, 1 subjects that would be used to create a representative functional atlas, and 0 otherwise.
◦ `out_dir`: Output directory location where statistical analysis results will be stored. This directory will be created if it does not exist.

**Figure 11:**
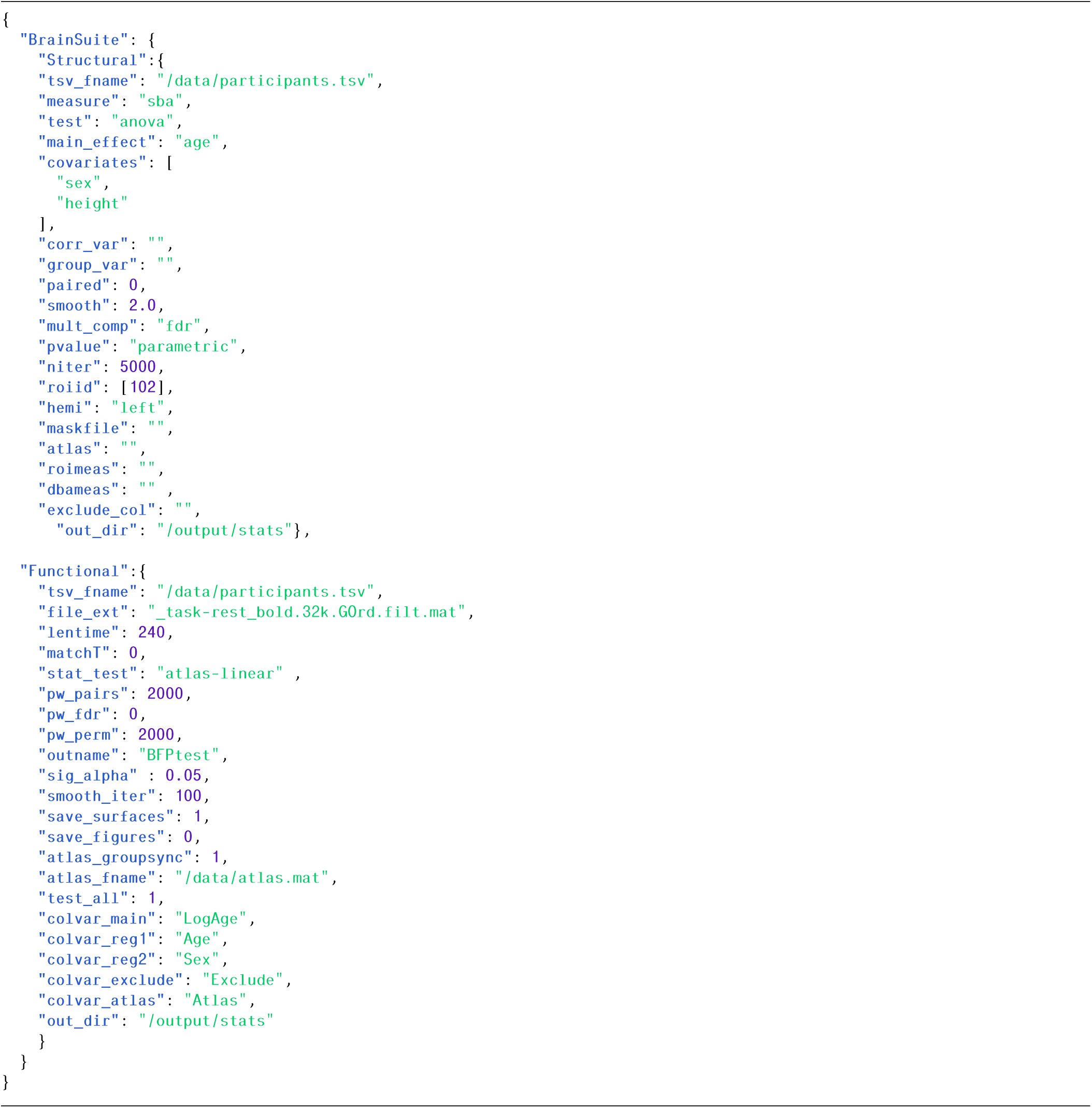
Statistical analysis model specifications.

## Footnotes

1 additional details about the BrainSuite atlases are available at https://brainsuite.org/atlases/.

2 details on the available FSL output types can be found here: https://fsl.fmrib.ox.ac.uk/fsl/fslwiki/FslEnvironmentVariables.

3 https://afni.nimh.nih.gov/pub/dist/doc/program_help/

